# Scalable flow synthesis of ultrasmall inorganic nanoparticles for biomedical applications *via* a confined impinging jet mixer

**DOI:** 10.1101/2025.11.12.688080

**Authors:** Andrea C. Kian, Mahima Gupta, Heeju Hong, Kálery La Luz Rivera, Nil Pandey, Katherine J. Mossburg, Derick N. Rosario-Berrios, Portia S. N. Maidment, Andrew R. Hanna, Priyash Singh, Zhenting Xiang, David Issadore, Andrew D. A. Maidment, Hyun Koo, David P. Cormode

## Abstract

Ultrasmall inorganic nanoparticles (sub-5 nm) have unique biomedical advantages due to rapid clearance, enhanced imaging contrast, and potent therapeutic properties. However, current synthesis methods are limited by low throughput, polydispersity, and reliance on harsh conditions such as organic solvents or high temperatures. We report a scalable, single-step aqueous synthesis using a confined impinging jet mixer (CIJM) that produces size-controlled, clinically relevant nanoparticles, including silver sulfide, silver telluride, cerium oxide, and iron oxide, under ambient conditions. The resulting nanoparticles are homogeneous, stable, and preserve their functional biological properties. We demonstrate consistent performance across scales, establishing the CIJM as a versatile and reproducible method for producing ultrasmall inorganic nanoparticles suitable for clinical translation and high-throughput biomedical applications.

## Introduction

Inorganic nanoparticles are versatile agents for biomedical imaging and therapy, with applications spanning magnetic resonance imaging (MRI), computed tomography (CT), photoacoustics, and targeted drug delivery.^1–4^ Ultrasmall nanoparticles (<5 nm) are of particular interest due to their ability to undergo renal clearance, reducing long-term organ retention and potential toxicity.^5–8^ Additionally, their high surface-to-volume ratios enhance catalytic activity and improved T_1_-weighted MRI contrast.^9–11^

Despite these advantages, current synthetic methods face major hurdles for clinical translation. Many require high temperatures, inert atmospheres, or undesirable organic solvents, limiting both scalability and biocompatibility.^12–14^ Conventional bulk synthesis approaches often yield polydisperse particles due to poor mixing control, leading to batch variability, low yields, and costly downstream processing, further reducing manufacturing efficiency and increasing production costs.^15–17^ There remains a critical need for scalable methods that consistently generate homogeneous nanoparticles.

Flow-based microfluidic systems offer improved control over reaction conditions and have been explored for nanoparticle synthesis.^14,18–20^ The confined impinging jet mixer (CIJM) stands out for its rapid mixing kinetics, clog-resistant design, low-cost scalability, and commercial availability (Fig. 1A).^21–23^ Although the CIJM has been deployed at industry scale for synthesizing mRNA-loaded lipid nanoparticles in COVID-19 vaccines,^24^ its potential for inorganic nanoparticle synthesis remains unexplored.

**Fig. 1.**
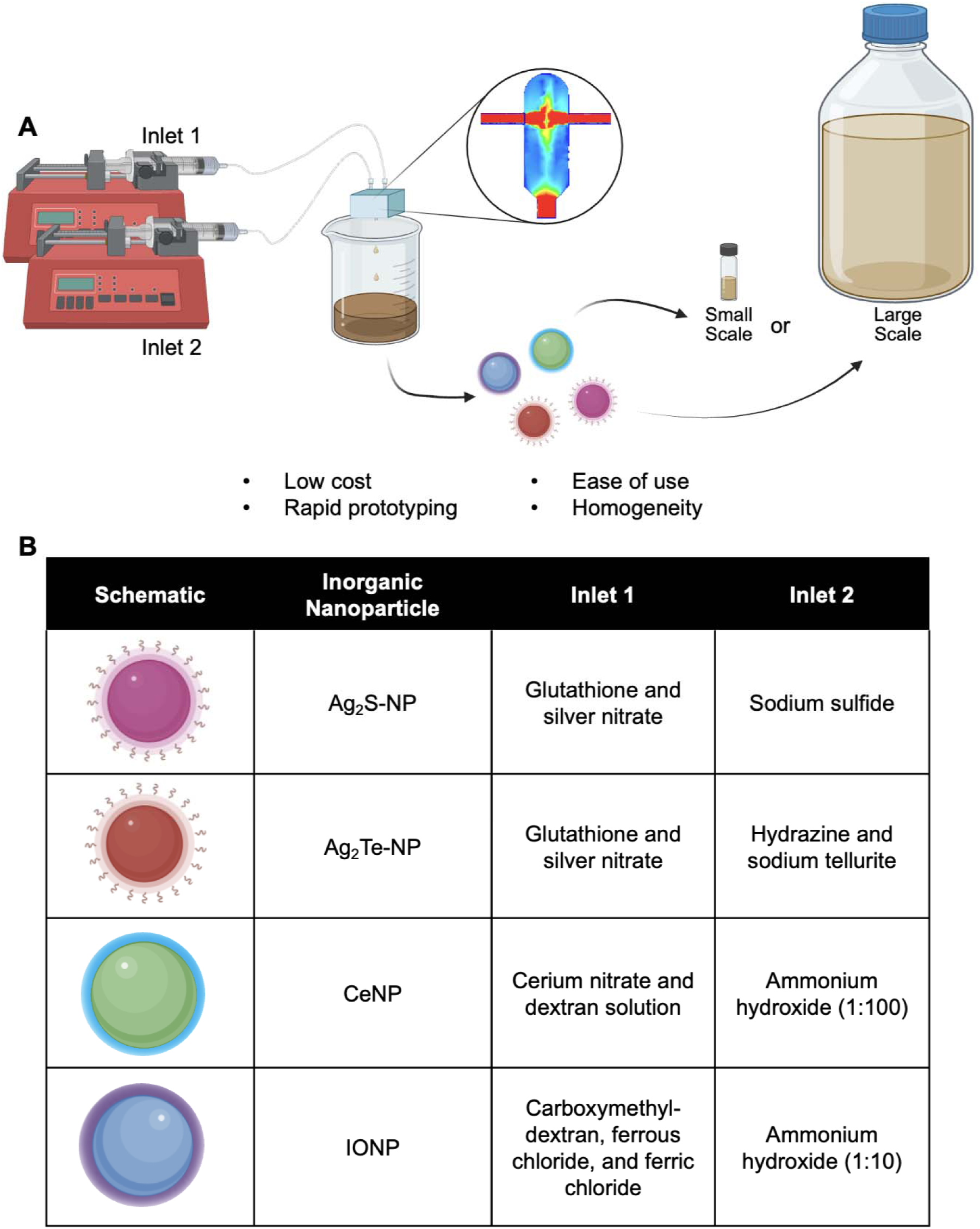
Confined impinging jet mixer (CIJM) for nanoparticle synthesis. A) Schematic of the CIJM showing the inlet streams and outlet collection point. Subset: Computational fluid dynamic (CFD) modelling of the CIJM highlighting turbulent mixing zone. B) Reagent pairs used at each inlet to synthesize Ag_2_S-NP, Ag_2_Te-NP, CeNP, and IONP *via* one-step aqueous-phase reactions.

Here, we evaluate the CIJM as a scalable platform for the one-step, aqueous-phase synthesis of four clinically relevant inorganic nanoparticles under ambient conditions: glutathione-coated silver sulfide (Ag₂S-NP), glutathione-coated silver telluride (Ag₂Te-NP), dextran-coated cerium oxide (CeNP), and carboxymethyl-dextran-coated iron oxide (IONP). These nanomaterials span a range diagnostic and therapeutic uses,^20,25–31^ including photoacoustic imaging, photothermal therapy, reactive oxygen species (ROS) scavenging, and antimicrobial biofilm disruption.^20,25,30–33^ IONP, in particular, are FDA-approved for treating iron deficiency,^34–36^ and have been repurposed clinically as an MRI contrast agent and an antibiofilm therapeutic.^26,29,37^

We demonstrate that the CIJM enables the facile synthesis of ultrasmall, stable, and homogeneous inorganic nanoparticles in aqueous solution, without organic solvents or elevated temperatures. The platform allows tunable core sizes and consistent performance across multiple applications, from biomedical imaging and catalytic protection from oxidative stress to biofilm degradation. Finally, we highlight the CIJM’s scalability by producing ∼1L of Ag₂S-NP in just 15 minutes, underscoring its potential for large-scale manufacturing in biomedical and translational settings.

## Results

To evaluate the versatility of the confined impinging jet mixer (CIJM) for inorganic nanoparticles synthesis, we prepared four formulations with distinct biomedical relevance: Ag₂S-NP, Ag₂Te-NP, CeNP, and IONP. These nanoparticles have been studied for applications in computed tomography (CT), contrast-enhanced mammography (CEM), photoacoustics, MRI, and catalytic therapeutics.^20,25–33,37^ The CIJM enables rapid, flow-based synthesis *via* micromixing at the collision point of two inlet streams (Fig. 1A). Reagents are delivered through each inlet, where turbulence and rapid mixing induces nanoprecipitation, and the resulting nanoparticles are collected immediately at the outlet. The method operates entirely in aqueous solutions at ambient temperature without the need for organic solvents. Each formulation was synthesized in a single step process using defined inlet chemistries (Fig. 1B), and the resulting products were reproducible and comparable to those obtained from established batch methods (Fig. S1).^21,38,39^ In addition to small-scale synthesis, we demonstrated scalability by producing ∼1L of Ag₂S-NP, confirming that nanoparticle quality was maintained under higher-throughput conditions.

### The CIJM Enables Synthesis of Ultrasmall, Size-Tunable Ag_2_S Nanoparticles

Silver sulfide nanoparticles (Ag_2_S-NP) synthesized using the CIJM exhibited ultrasmall core sizes, as confirmed by transmission electron microscopy (TEM), UV/visible absorbance spectra (UV/Vis), and X-ray photoelectron spectroscopy (XPS) consistent with previous reports (Fig. 2A-C and Fig. S2).^40,41^ We assessed nanoparticle stability in biologically relevant media (H_2_O, PBS, and PBS + 10% FBS) and observed no changes over seven days, indicating that CIJM-produced Ag_2_S-NP remain stable under physiological conditions (Fig. S3).

**Fig. 2.**
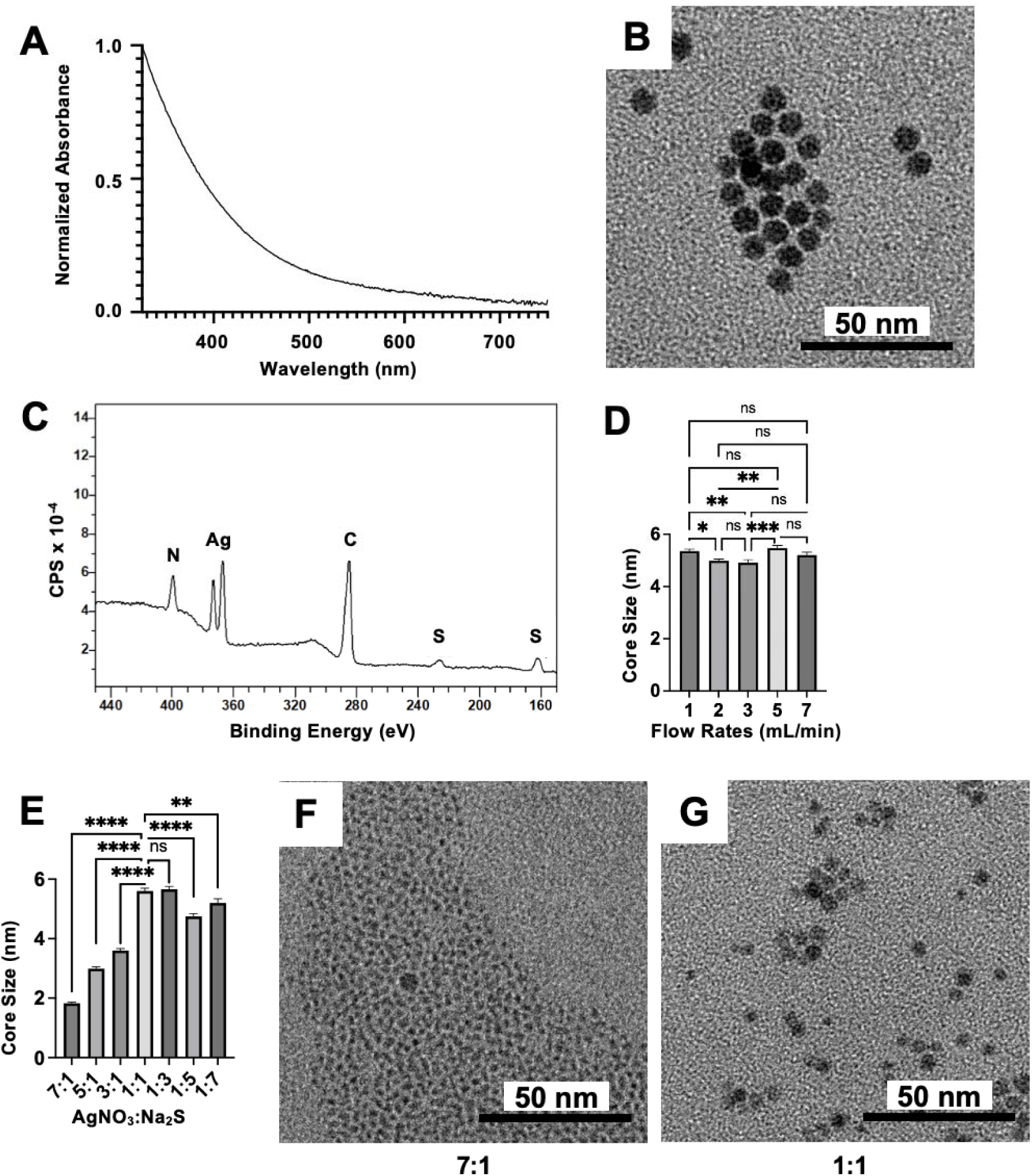
Characterization and synthesis parameters of Ag_2_S-NP. A) UV/Vis absorption spectra of Ag_2_S-NP. B) TEM images of Ag_2_S-NP. C) XPS spectra of Ag_2_S-NP. D) Core sizes of Ag_2_S-NP synthesized at different flow rates (1, 2, 3, 5, and 7 mL/min). E) Core sizes of Ag_2_S-NP synthesized at different flow rate ratios. F-G) TEM of selected Ag_2_S-NP prepared using different flow rate ratios. Scale bars = 50 nm. ns = non-significant, * p≤0.05, ** p≤0.01, *** p≤0.001, **** p≤0.0001. Error bars represent the standard error of the mean.

To evaluate tunability of particle size, a key parameter for physical properties and pharmacokinetics, we investigated three synthesis parameters: total flow rate, flow rate ratio (AgNO_3_:Na_2_S), and reagent concentration. Flow rate, varied from 1-7 mL/min per inlet, had minimal effect on particle size, yielding consistent nanoparticles of ∼5.2 ± 0.1 nm across conditions (Fig. 2D, Fig. S4A). Although size was unaffected, higher flow rates increased production throughput, consistent with observations in organic nanoparticle systems where core size plateaus beyond a critical Reynolds number.^42,43^ A maximum rate of 7 mL/min/inlet was used, reflecting the upper limit of our syringe pumps.

In contrast, the flow rate ratio had a pronounced effect. At a 7:1 AgNO₃:Na₂S ratio, nanoparticles averaged 1.8 nm, while more balanced ratios produced progressively larger cores, plateauing at ∼5.3 nm at 1:1 (Fig. 2E–G, Fig. S4B). Thus, flow rate ratio is a robust parameter for tuning the Ag_2_S-NP size within the sub-5 nm range. By comparison, reagent concentration (increased up to 30x) had no measurable effect on core size (Fig. S5), indicating that nanoparticle yields can be scaled without altering physical properties. Together, these results establish the CIJM as a tunable and scalable platform for producing ultrasmall Ag₂S-NPs under ambient conditions.

### The CIJM Enables Synthesis of Ultrasmall, size tunable Ag₂Te Nanoparticles

Silver telluride nanoparticles (Ag₂Te-NP), analogous to their silver chalcogenide counterpart Ag₂S-NP, were successfully synthesized using the CIJM. In this approach, hydrazine was added to a mixture of silver nitrate and sodium tellurite to generate Te²⁻ *in situ*, enabling nanoparticle formation.^44,45^ The resulting Ag₂Te-NP were stable, with a mean core diameter size of 2.1 nm as determined by TEM, UV/Vis, and XPS (Fig. S6-8). Flow rates (1 to 7 mL/min per inlet) and flow rate ratio variations had negligible effects on core size, consistent with Ag_2_S-NPs (Fig. S9 and S10). In contrast, increasing hydrazine concentration to 1.6 M produced larger particles (2.7 ± 0.2 nm versus ∼2.1 nm for lower concentrations), demonstrating that reducing agent strength can modulate nanoparticle growth (Fig. 3A-E).

**Fig. 3.**
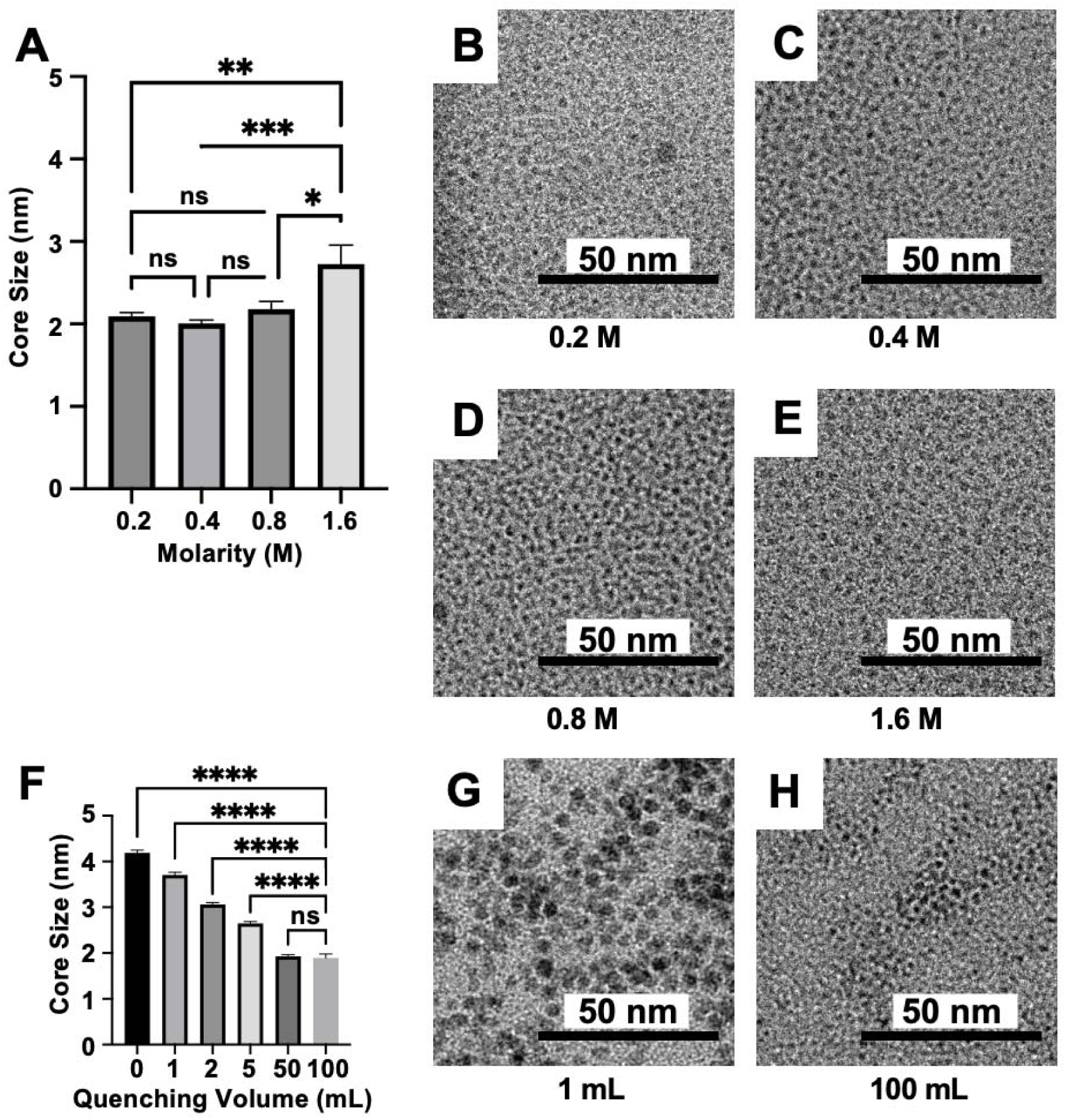
Size-tuning parameters for Ag_2_Te-NP. A) Core sizes of Ag_2_Te-NP synthesized using different hydrazine concentrations (*i.e.* 0.2, 0.4, 0.8, 1.6 M). B-E) TEM images of Ag_2_Te-NP synthesized at the indicated hydrazine concentrations. F) Core sizes of Ag_2_Te-NP synthesized with different quench volumes (*i.e.* 0, 1, 2, 5, 50, 100 mL). G-H) TEM images of Ag_2_Te-NP synthesized at 1 and 100 mL quench volumes. Scale bars = 50 nm. ns = non-significant, * p≤0.05, ** p≤0.01, *** p≤0.001, **** p≤0.0001. Error bars represent the standard error of the mean.

To further evaluate tunability, we varied the quenching volume (defined as the volume of DI water added post-synthesis). Increasing quenching volume led to progressively smaller nanoparticles, with mean diameters decreasing from 4.2 ± 0.1 nm (no quenching) to 1.9 ± 0.1 nm (100 mL DI water, Fig. 3F-H and Fig. S11). This trend suggests that dilution suppresses nanoparticle growth, consistent with previous reports in IONP systems.^46,47^ Together, these findings demonstrate the CIJM’s compatibility with Ag₂Te-NP synthesis and identify two tunable parameters, reducing agent concentration and quench volume, that enable control over core size.

### The CIJM Enables ambient synthesis of CeNP with limited size tunability

Cerium oxide nanoparticles (CeNP) were synthesized in the CIJM by mixing an ammonia solution and a cerium nitrate/dextran solution, where the resulting pH decrease induced nanoparticle formation.^28,48^ Initial attempts using reagent concentrations from prior protocols^28,49,50^ caused mixer clogging; however, dilution of both streams prevented blockage while still producing nanoparticles (Fig. S12). Notably, the CIJM method required no heating, in contrast to conventional synthesis approaches.

Stable nanoparticle formation was confirmed by UV/Vis, TEM, and XPS, which showed a characteristic absorbance peak at 290 nm, a mean core size of 2 nm, and the elemental peaks consistent with cerium (Fig. S13 and S14). These nanoparticles were smaller than those produced by batch methods (3.5 nm).^50^ To evaluate size tunability, we varied total flow rate, flow rate ratio, and post-synthesis mixing duration. Across all tested conditions, only modest or negligible changes in core size were observed (Fig. 4A–I, Fig. S15–S18), indicating that, unlike Ag-based NP systems, CeNP synthesis *via* CIJM offers limited control over particle size with the current parameters. Nonetheless, the platform supports reproducible, ambient-condition synthesis of ultrasmall CeNPs.

**Fig. 4.**
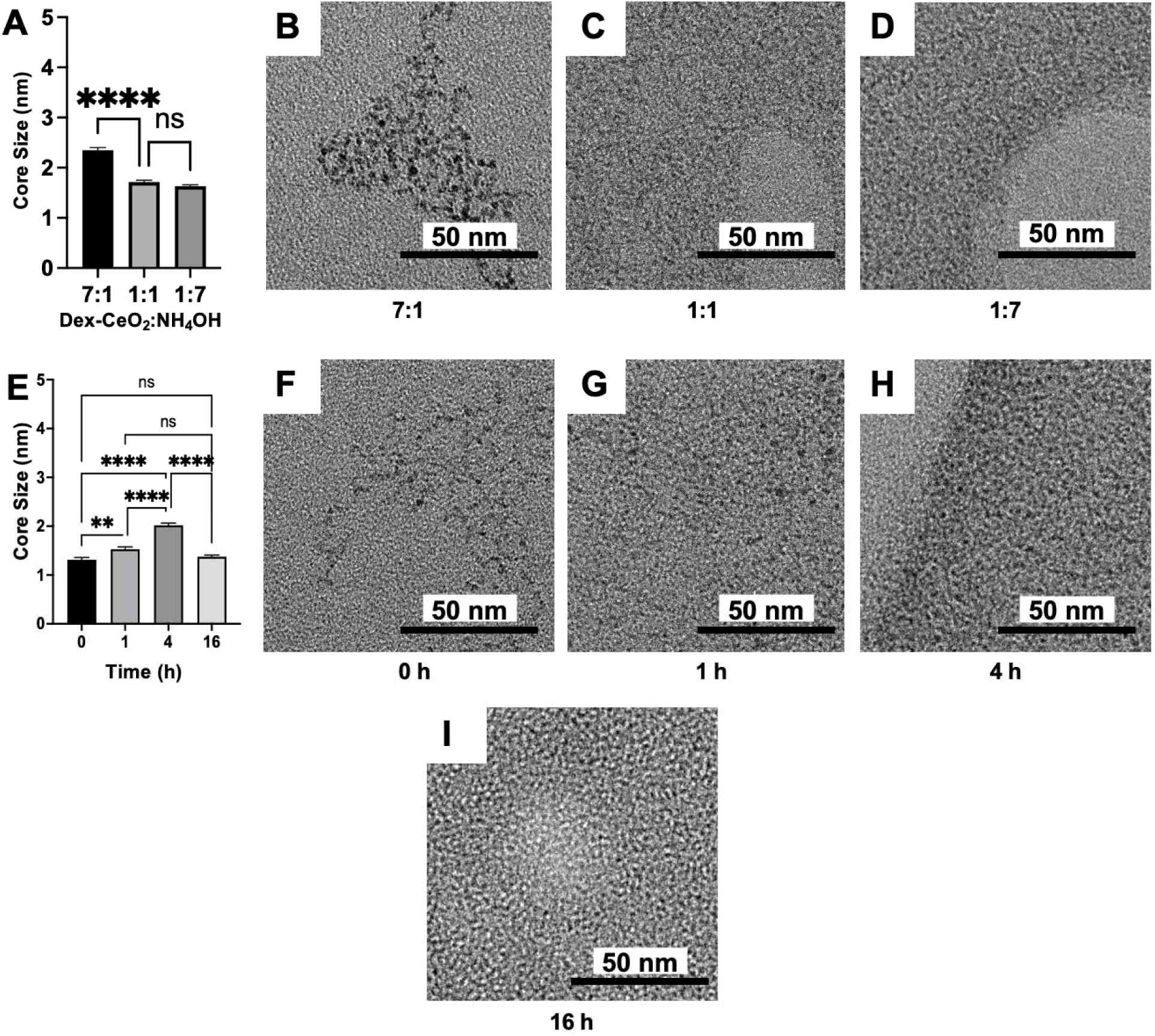
**CeNP synthesis parameters**. A) Core sizes of CeNP synthesized at different flow rate ratios (7:1, 1:1, and 1:7). B-D) TEM images of CeNP prepared at the indicated ratios. E) Core sizes of CeNP synthesized after varying post-synthesis mixing times (0, 1, 4, and 16 hours). F-I) TEM images of CeNP from the corresponding mixing times. Scale bars = 50 nm. ns = non-significant, * p≤0.05, ** p≤0.01, *** p≤0.001, **** p≤0.0001. Error bars represent the standard error of the mean.

### The CIJM Enables CMD-coated IONP Synthesis with selective size tunability

Iron oxide nanoparticles (IONP) were synthesized by introducing ammonium hydroxide (NH_4_OH) into a solution of iron salts and carboxymethyl dextran (CMD) within the CIJM. Compared to conventional bulk synthesis, which typically requires heating to 90 °C for one hour and an inert atmosphere,^51,52^ the CIJM approach was conducted at room temperature and under ambient conditions, significantly reducing processing time and complexity.

Stable IONP formation was confirmed by UV/Vis, TEM, and XPS (Fig. S19 and S20). We next investigated tunability of core size and hydrodynamic diameter, as well as their catalytic activity (Fig. S21-24). Core size was largely unaffected by flow rate (1 to 7 mL/min/inlet) or flow rate ratio (3:1 to 1:7), with particles averaging 2.2-2.3 ± 0.1 nm (Fig. S21 and S22). Increasing CMD concentration from 5 to 350mg moderately increased core diameter, with 300mg yielding IONP of 2.90±00.10nm (Fig. 5A–D). By contrast, varying NH₄OH concentration (0.7–11.2%) strongly influenced core size, producing particles as small as 1.70nm at 0.7% NH₄OH (Fig. 5E–H). These results indicate that ligand and base concentrations, but not flow conditions, provide control over IONP core size. Catalytic activity of the formulations was comparable, consistent with their similar core sizes (Fig. S23).

**Fig. 5.**
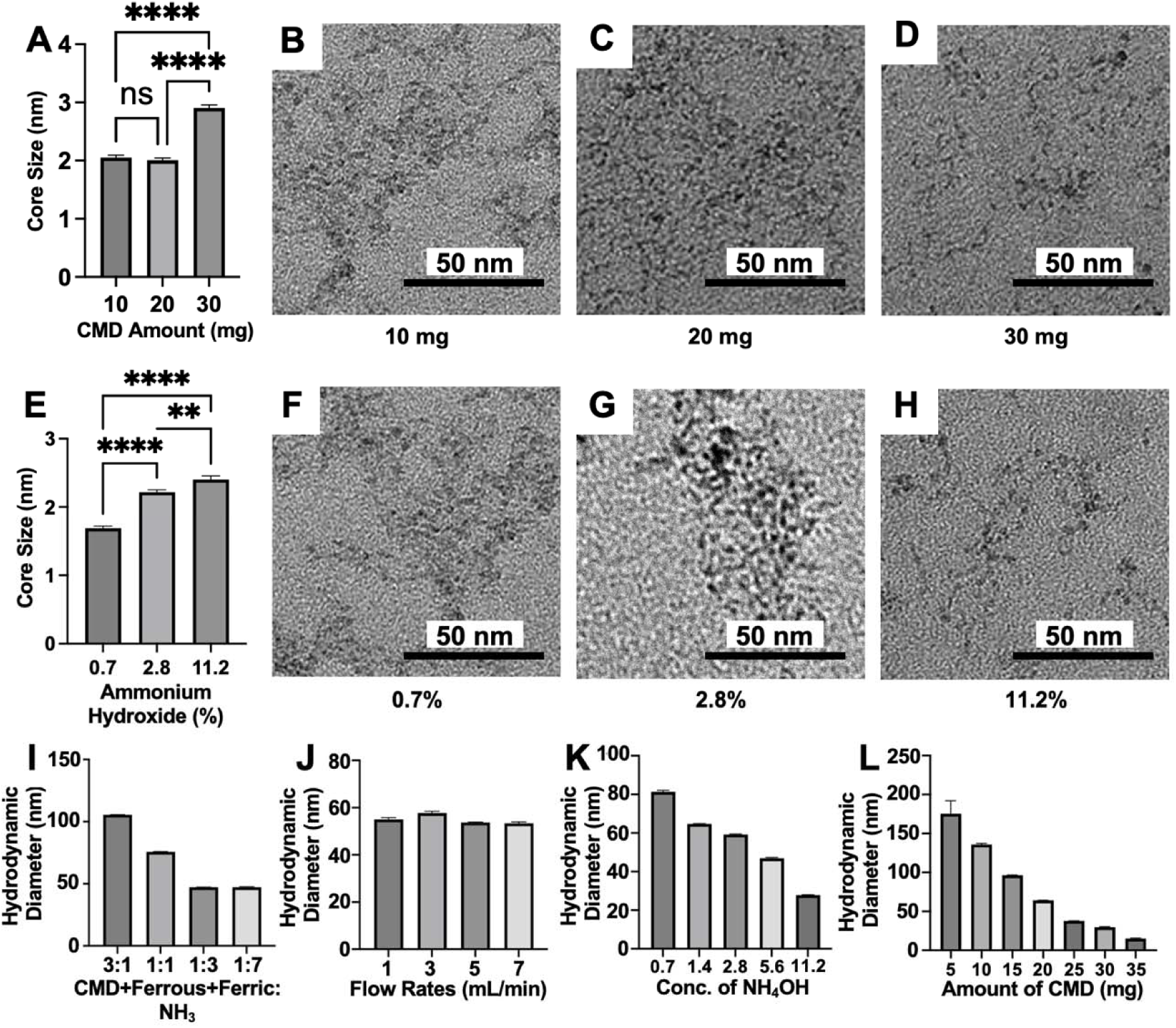
Parameters influencing IONP size. A) Core sizes of IONP synthesized with different CMD concentrations (10, 20, 30 mg). B-D) TEM images of IONP synthesized at the indicated CMD amounts. E) Core sizes of IONP synthesized with varying NH_4_OH concentrations (0.7, 2.8, and 11.2%). F-H) TEM images of IONP synthesized at the indicated NH_4_OH concentrations. I) Hydrodynamic diameters of IONP synthesized at different flow rate ratios (3:1 to 1:7). J) Hydrodynamic diameters at different total flow rates. K) Hydrodynamic diameters at varying NH_4_OH concentrations (0.7 to 11.2%). L) Hydrodynamic diameters at different amounts of CMD (5 to 35 mg). Scale bars = 50 nm. ns = non-significant, * p≤0.05, ** p≤0.01, *** p≤0.001, **** p≤0.0001. Error bars represent the standard error of the mean.

Notably, hydrodynamic diameter was tunable across a broad range. Adjusting flow rate ratio from 1:3 to 3:1 increased hydrodynamic size from 47.2 to 105.5 nm (Fig. 5I). whereas absolute flow rate (1 to 7 mL/min) had no effect (Fig. 5J). NH_4_OH concentration inversely modulated hydrodynamic diameter, decreasing from from 81.3 to 27.8 nm as concentration rose from 0.7 to 11.2% (Fig. 5K). Counterintuitively, increasing CMD from 5 to 35 mg reduced hydrodynamic diameter from 175.2 to 15.1 nm (Fig. 5L). These findings demonstrate that, while IONP core size is only modestly tunable, hydrodynamic diameter, a key property for colloidal stability and *in vivo* performance, can be effectively controlled during CIJM-based synthesis.

### CIJM-synthesized nanoparticles are functional in exemplar biomedical applications

To assess the biomedical relevance of CIJM-synthesized nanoparticles, we investigated their performance in experimental models specific to each type.

Ag₂S-NP and Ag₂Te-NP were evaluated for x-ray imaging using contrast-enhanced mammography (CEM) and computed tomography (CT) phantoms.^25,30,31,33,53^ At equivalent silver concentrations, both nanoparticle formulations outperformed iopamidol (a clinical iodinated contrast agent) and their corresponding metal salts. In CEM imaging, Ag₂Te-NP and Ag₂S-NP exhibited the highest contrast-to-noise ratios (CNRs) with CNRs of 19 and 10, respectively, compared to ∼9 for tellurium and silver salts and ∼6 for iopamidol (Fig. 6A–B, Fig. S24). Ag_2_Te-NP generated higher contrast than Ag_2_S-NP, attributable to tellurium’s imaging properties. In CT phantom imaging, attenuation rates were 76 HU·mg/mL (Ag_2_Te-NP), 61 HU·mg/mL (Ag_2_S-NP), and 46 HU·mg/mL (iopamidol), confirming superior contrast enhancement compared to the clinical standard (Fig. 6C-D).

**Fig. 6.**
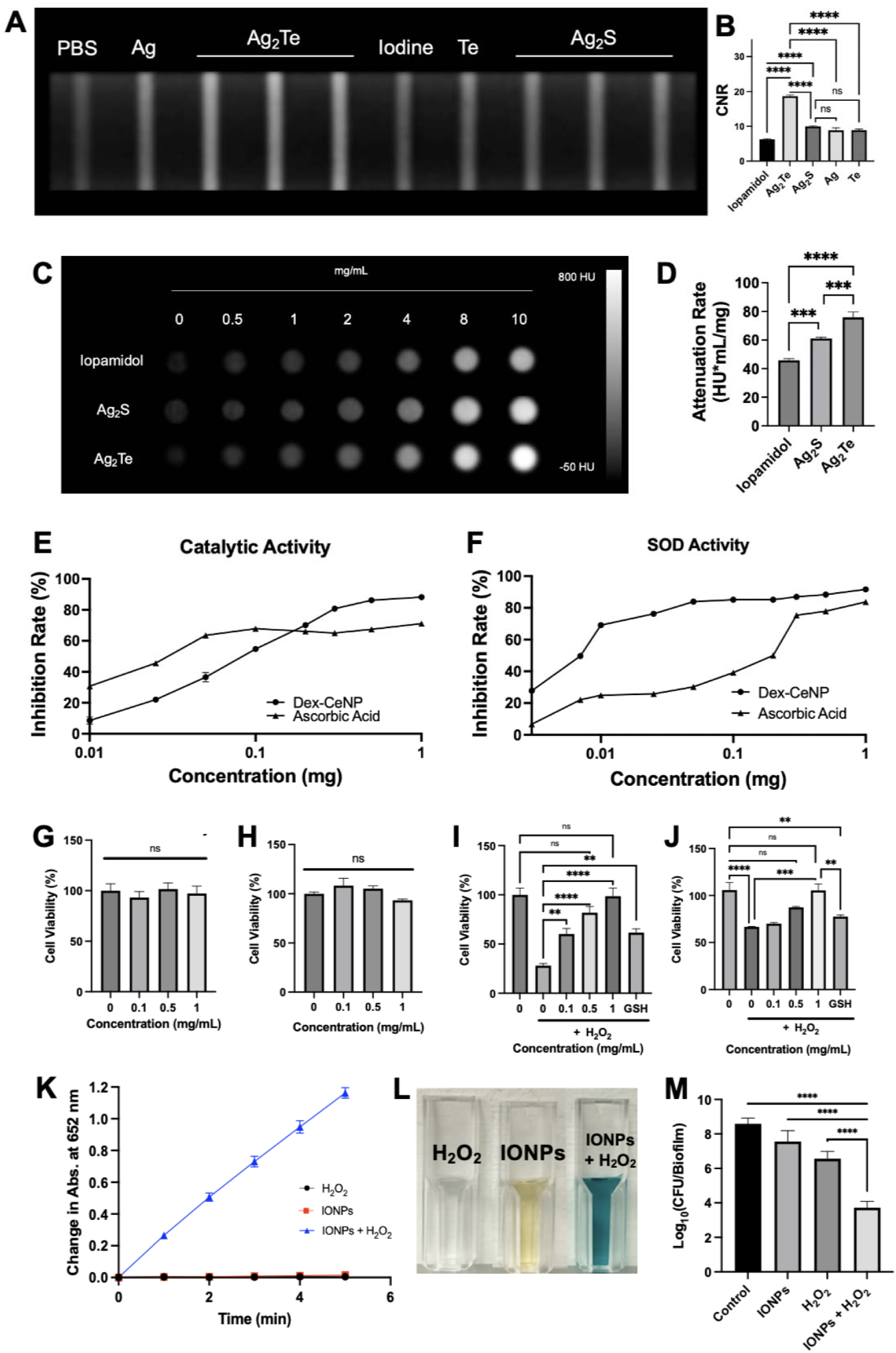
**Functional activity of CIJM-synthesized nanoparticles**. A) CEM images of Ag_2_S-NP and Ag_2_Te-NP. B) Corresponding contrast-to-noise ratios (CNR). C) CT phantom images of Ag_2_S-NP and Ag_2_Te-NP. D) Attenuation rates derived from CT scans. E-F) CAT- and SOD-mimetic activity of CeNP compared to ascorbic acid. G-J) Cell viability and ROS-scavenging activity of CeNP in BJ5ta and C2bbe1 cells exposed to H_2_O_2_. K) Change in absorbance of TMB at 652 nm for IONP with H_2_O_2_, IONP alone, and H_2_O_2_ alone. L) Photograph of samples from K. M) Antibacterial activity of IONP with H_2_O_2_ as determined by colony-forming units (CFU). ns = non-significant, * p≤0.05, ** p≤0.01, *** p≤0.001, **** p≤0.0001. Error bars represent the standard error of the mean.

CeNP were tested for antioxidant capacity. Both catalase (CAT)- and superoxide dismutase (SOD)-like activities were detected in CIJM-synthesized CeNP (Fig. 6E–F). To assess cellular protection from oxidative damage, ^28,50^ fibroblast (BJ5ta) and colorectal epithelial (C2BBe1) cells were exposed to hydrogen peroxide (H_2_O_2_) with or without CeNP co-incubation.

^27^ While CeNP alone were non-toxic, H_2_O_2_ exposure reduced cell viability by 72% (BJ5ta) and 33% (C2BBe1). Co-treatment with CeNP and H_2_O_2_ restored viability to baseline levels, confirming their protective, antioxidant function (Fig. 6G–J).

IONP were evaluated for catalytic (peroxidase-like) and antimicrobial/antibiofilm activities.^26,34–37^ In a TMB colorimetric assay,^26,54,55^ IONP combined with H_2_O_2_ induced rapid substrate oxidation, demonstrating peroxidase-like activity (Fig. 6K–L). In parallel, CIJM-synthesized IONP effectively disrupted *Streptococcus mutans* biofilms, killing 99.9% of bacterial cells within 5 minutes when combined with H_2_O_2_, a >500-fold increase in biocidal efficacy compared to either treatment alone (Fig. 6M).

Together, these findings validate that CIJM-synthesized nanoparticles not only match but in some cases exceed the functional performance of conventionally prepared analogs, while offering rapid, scalable, and ambient-condition synthesis without harsh solvents.

### CIJM enables scalable and consistent nanoparticle synthesis

To evaluate the scalability of the CIJM platform, we performed a large-scale synthesis of Ag_2_S-NP. Scale-up was achieved by increasing both the flow rate (from 7 to 30 mL/min per inlet) and the total reagent volume (100-fold increase), resulting in a theoretical 100-fold increase in nanoparticles output within minutes (Fig. 7A). Inductively coupled plasma optical emission spectroscopy (ICP-OES) confirmed a ∼78-fold increase in total silver yield, from 1.1 to 86.1 mg (Fig. 7B). Nanoparticles from small- and large-scale batches were indistinguishable in core size and spectral properties, as verified by TEM and UV/Vis (Fig. 7C-E and Fig. S25), indicating that scale-up did not affect product quality.

**Fig. 7.**
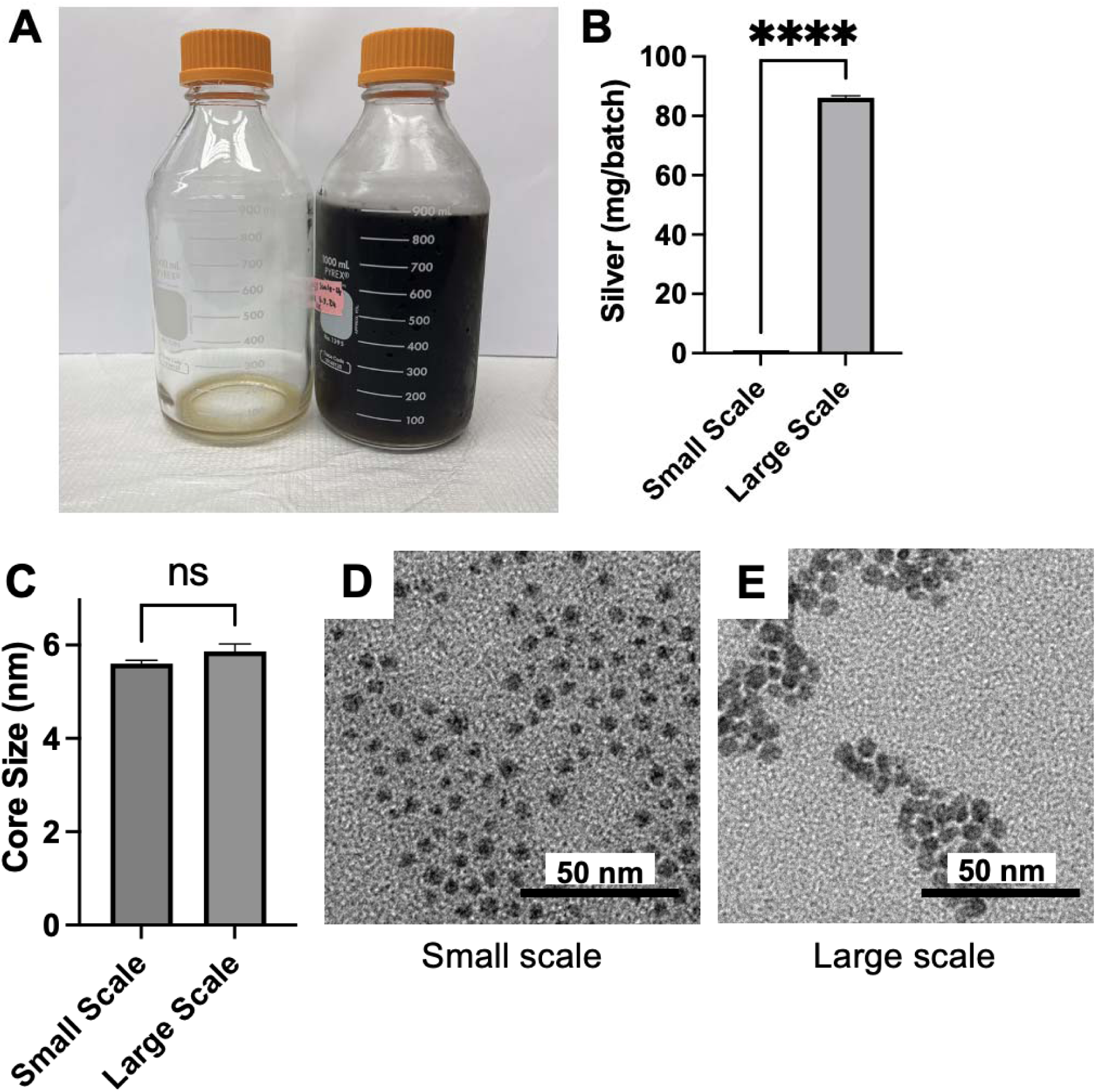
Scale up of Ag2S-NP with the CIJM. A) Photograph of Ag_2_S-NP synthesized at small scale (left) and after scale-up (right). B) Quantification of nanoparticle yield. C) Core sizes of small- and large-scale nanoparticles. D-E) TEM images of the nanoparticles. Scale bars = 50 nm. ns = non-significant, * p≤0.05, ** p≤0.01, *** p≤0.001, **** p≤0.0001. Error bars represent the standard error of the mean.

These results demonstrate that the CIJM supports high-throughput production of ultrasmall inorganic nanoparticles without compromising homogeneity or stability, enabling consistent synthesis across scales. Collectively, our findings establish the CIJM as a reproducible, aqueous-phase platform for synthesizing diverse ultrasmall inorganic nanoparticles with tunable or controllable sizes, preserved functional activity, and scalability to production-relevant volumes. By operating entirely under ambient conditions, without organic solvents, elevated temperatures, or inert atmospheres, the CIJM provides a versatile and generalizable platform for streamlined development of clinically relevant nanomaterials.

## Discussion

This study demonstrates that the confined impinging jet mixer (CIJM) enables rapid, reproducible, and solvent-free synthesis of multiple ultrasmall inorganic nanoparticles, including silver sulfide (Ag₂S), silver telluride (Ag₂Te), cerium oxide (CeNP), and iron oxide (IONP), under ambient conditions. The nanoparticles produced were homogeneous in size and morphology, as confirmed by TEM, UV/vis, and XPS, and several formulations exhibited tunable core diameters through adjustments to flow rate ratios, reagent concentrations, or quenching conditions. While originally developed for organic nanoparticle production, our results establish the CIJM as a scalable and cost-effective platform for inorganic nanoparticle synthesis. Notably, tunability varied by material: Ag₂S-NP size was highly responsive to flow rate ratios, Ag₂Te-NP were more sensitive to reducing agent and quench conditions, and CeNP and IONP responded to ligand concentration or base strength. These findings suggest that CIJM-mediated nanoparticle synthesis depends on both particle-specific chemistries and microfluidic parameters, offering opportunities for targeted optimization.

The CIJM-synthesized nanoparticles retained hallmark functional properties relevant to biomedical applications. Ag₂S-NP and Ag₂Te-NP outperformed iodinated contrast agents (the clinical standard) in CT and CEM phantom imaging. CeNP displayed strong CAT- and SOD-like activity and protected mammalian cells from oxidative stress. IONP demonstrated peroxidase-like activity and effectively eradicated cariogenic biofilms in acidic pH environment. These results collectively validate that CIJM-synthesized inorganic nanoparticles are not only structurally well-formed but also biologically active and application-ready.

We further showed that Ag₂S-NP synthesis could be scaled up ∼100-fold, generating a large batch within 15 minutes without compromising nanoparticle quality. TEM and spectroscopic analyses confirmed equivalence between small- and large-scale batches. While other microfluidic approaches have achieved high-volume synthesis, they often require custom fabrication or complex infrastructure. In contrast, the CIJM offers a commercially available, low-cost, and scalable solution that can be rapidly implemented for industrial or clinical nanoparticle production.

Previous work has shown that devices such as herringbone, T-junction, and Y-junction mixers can be used for inorganic nanoparticle synthesis, and more advanced chaotic mixers can achieve wide size ranges (nm to µm) with high reproducibility.^56–59^ Parallel herringbone devices, in particular, can produce higher yields than those reported here by employing hundreds of parallel channels.^20,59^ However, the CIJM uniquely combines rapid micromixing, clog resistance, and scalability in a simple, commercially available format without the need of complex flow arrangements. These features facilitate cost-effective industrial implementation of clinically-relevant nanoparticle doses. Given its demonstrated use in producing polymeric and lipid nanoparticles with core sizes >100 nm,^43,60^ the CIJM could be adapted for a broader range of inorganic nanomaterials, including larger constructs or those requiring more complex ligand chemistries.

While promising, our study has limitations. First, experiments explored a limited flow rate range (1–300mL/min/inlet), and CIJM performance at higher pressures remains uncharacterized. Second, while the 1 to 60nm size range is well-suited for renal clearance and biomedical use, some applications, such as nanorobotics, may require larger particles.^61–65^ Third, all biological validation was performed in benchtop models; while these results are highly supportive of the nanoparticles’ therapeutic potential, *in vivo* studies will be required to confirm pharmacokinetics, biodistribution, and efficacy.

## Conclusions

Overall, this work establishes the CIJM as a versatile and generalizable platform for the rapid, reproducible, scalable, and tunable synthesis of diverse ultrasmall inorganic nanoparticles under ambient, solvent-free conditions. By enabling precise control over particle formation across multiple chemistries, the CIJM bridges a key gap between laboratory-scale discovery and clinically relevant manufacturing. The resulting nanoparticles retain hallmark structural integrity and functional activity, with early biological validation highlighting therapeutic potential in imaging, oxidative stress protection, and antimicrobial applications. Together, these findings position the CIJM as a broadly enabling technology for the streamlined development of clinically relevant nanomaterials, paving the way for future studies to expand its use to additional inorganic systems, explore higher-throughput configurations, and validate *in vivo* performance for clinical implementation.

## Materials and Methods Materials

Silver nitrate (AgNO_3_, 99%), sodium sulfide (Na_2_S), L-glutathione (GSH, 98%), sodium tellurite (Na_2_TeO_3_, 97%), cerium nitrate hexahydrate (Ce(NO_3_)_3_, 99.99%), ferrous chloride tetrahydrate (FeCl_2_•4H_2_O, 98%), ferric chloride hexahydrate (FeCl_3_•6H_2_O, 99.99%), and carboxymethyl-dextran were all purchased from Sigma Aldrich (St. Louis, MO). Sodium hydroxide (NaOH), hydrazine hydrate (N_2_H_4_, 100%), nitric acid (HNO_3_), and ammonium hydroxide (NH_4_OH, 28-30%) were purchased from Fisher Scientific (Pittsburgh, PA). Saliva-coated hydroxyapatite disks were purchased from Clarkson Chromatography Products Inc. (South Williamsport, PA). Dextran T-10 was purchased from Pharmacosmos (Holbaek, Denmark). Iopamidol (ISOVUE-300) was purchased from Bracco Diagnostics (Monroe Township, NJ). A confined impinging jet mixer (CIJM) was purchased from Holland Applied Technologies (Burr Ridge, IL).

## Methods

### Confined Impinging Jet Mixer

In brief, syntheses were performed using the confined impinging jet mixer (CIJM) by flowing through it two different aqueous solutions simultaneously (Fig. 1B). Braintree Scientific syringe pumps (BS-300) were used to inject the solutions into the mixer at the desired flow rate. Within the mixer, internal micromixing occurs, and the product is collected in a flask at the outlet. In order to illustrate this micromixing process, previously reported values were used to model the CIJM using Autodesk Fusion and Computation Fluid Dynamics (Fig. 1A).^38,66^

## Nanoparticle Synthesis

### GSH-Coated Sub-5 nm Silver Sulfide Nanoparticles

The method for synthesizing sub-5 nm, Ag_2_S-NP was adapted from a previous study.^25^ An aqueous solution (5 mL) containing 2.8 mg AgNO_3_ and 52 mg GSH (pH 7.4) was placed into a syringe connected to inlet one. Another aqueous solution (5 mL) containing 2 mg Na_2_S was placed into a syringe connected to inlet two. A flow rate of 7 mL/min/inlet was used for the syntheses (unless otherwise specified), and the resulting product was stirred in ambient conditions for 24 hours. The nanoparticles (NP) were thereafter washed thrice at 4000 rpm centrifugation using DI water in 3 kDa MWCO tubes and then concentrated to 1 mL. The NP were filtered using 0.02 µm filters and stored at 4 °C until further use.

The scaled-up Ag_2_S-NP solution was prepared according to the same protocol above, using 100x more solution for each inlet. The two solutions were placed in pressurized vessels (5-gallon and 3-gallon stainless steel vessels) connected to a nitrogen tank with a pressure controller regulating the flow rate.^67^ For this study, the minimum pressure of 2 psi generated a flow rate of approximately 30 mL/min/inlet. The collected output then followed the previous protocol for washing and filtering.

### GSH-Coated Sub-5 nm Silver Telluride Nanoparticles

A sub-5 nm core, Ag_2_Te-NP were synthesized using a method adapted from a previous publication.^30^ The aqueous solution (8 mL), containing 10 mg AgNO_3_ and 83 mg GSH was loaded into a syringe connected to inlet one. Another aqueous solution (8 mL), containing 15.3 mg Na_2_TeO_3_ and 1 mL of N_2_H_4_ was connected to inlet two. The flow rate of 7 mL/min/inlet was used for the synthesis (unless otherwise specified). The reaction was collected into a flask containing 100 mL of DI water and then stirred for 5 minutes at room temperature. The NP were then washed thrice by 4000 rpm centrifugation in 3 kDa molecular weight cut-off tubes (MWCO) using DI water and concentrated to 1 mL. The NP were filtered using 0.45 µm filters and stored at 4 °C until further use.

### Dextran-Coated Sub-5 nm Cerium Oxide Nanoparticles

The method for synthesizing sub-5 nm, CeNP was adapted from a previous study.^28,50^ An aqueous solution (6 mL) containing 20 mg of T-10 dextran and 8.6 mg of Ce(NO_3_)_3_ was loaded into a syringe connected to inlet one. Another aqueous solution (6 mL), containing 1% NH_4_OH was connected to inlet two. The flow rate of 7 mL/min/inlet was used for the synthesis (unless otherwise specified). The reaction was collected in a flask and stirred overnight. The NP were centrifuged at 4000 rpm for 10 minutes to remove any large aggregates. The supernatant was then transferred to a 3 kDa MWCO and washed six times using DI water and concentrated to 1 mL. The NP were filtered using 0.45 µm filters and stored at 4 °C until further use.

### Carboxymethylated Dextran-Coated Sub-5 nm Iron Oxide Nanoparticles

The protocol for synthesizing IONP was devised for this study. A vial containing 1 mL of DI water and 20 mg of CMD was stirred for five minutes. In a separate vial, 7.2 mg of FeCl_2_ 4H_2_O and 14.7 mg of FeCl_3_ 6H_2_O were dissolved in 0.5 mL of DI water. This solution was then added to the CMD solution. The resulting mixture was stirred for five minutes. In the CIJM, the syringe at inlet one contained 1.5 mL of 2.8% ammonium hydroxide and the other contained 1.5 mL of the CMD-ferrous-ferric solution. The solutions were injected into the CIJM at 7 mL/min/inlet, unless otherwise specified. The mixture was then stirred for two hours. The NP were washed with DI water in 100 kDa MWCO tubes five times under 4000 rpm centrifugation and then concentrated to 1 mL. The NP was then stored at 4 °C until further use.

## Nanoparticle Characterization

### UV/Vis Spectrometry

A Genesys 150 UV/Visible Spectrophotometer (Thermo Scientific) recorded the UV/Visible spectra. The samples were prepared by diluting the samples to 10 ug/mL and placing 1 mL into a cuvette for analysis. On graphs, the absorption intensity maximum was normalized to one for all samples.

### Dynamic Light Scattering

A Malvern Zetasizer was used to measure the particle size of IONP. Samples were prepared by diluting 100 µL of the nanoparticles into 900 µL of DI water and then centrifuged at 14.1 RCF. The supernatant was collected and measured with DLS.

### Transmission electron microscopy

A Tecnai T12 electron microscope, which operates at 100 kV, was used to obtain TEM micrographs for core size measurements. The samples were prepared by placing 10 µL of the concentrated nanoparticles onto a carbon-coated copper grid with 200 mesh (Electron Microscopy Science). ImageJ (National Institutes of Health, Bethesda, MD) was used to measure the core sizes of 100 nanoparticles per sample. Comparison of all core sizes from the differing synthesis parameters used a one-way ANOVA with Tukey’s multiple comparisons test.

### X-ray photoelectron spectrometer

A Physical Electronics VersaProbe 5000 X-ray photoelectron spectrometer with a monochromatic Al K-Alpha source of 1486.2 eV was used to analyze the elemental composition of the nanoparticles. Samples were prepared by lyophilizing 1 mL of solution containing 10 mg of NP. The samples were loaded onto a 2 inch round sample holder and samples were acquired using a 200 micron 50 W electron beam. The pass energy for the survey and high-resolution spectra was set to 117 and 27 eV, respectively. Dual source neutralization with a combination of ion gun and electron neutralization was used during analysis. The data was charge corrected to adventitious carbon of 284.8 eV binding energy.

### Inductively-coupled plasma optical emission spectroscopy

A Spectro Genesis ICP-OES was used to determine silver, cerium, or iron content depending on the sample being analyzed. The samples were prepared by adding 10 µL of the concentrated NP solution to 1 mL of nitric acid, overnight. 9 mL of DI water was then added.

### Phantom Imaging

#### Contrast-Enhanced Mammography

A contrast-embedded 5 cm thick tissue-equivalent phantom was used to examine the contrast properties of the silver nanoparticles.^25^ The center portion of the phantom varied linearly from 100% glandular to 100% adipose tissue equivalent. Adipose equivalent layers were added to the top and bottom of the phantom simulated the skin. Samples were prepared at 10 mg/mL (in terms of silver or iodine as appropriate) of silver sulfide, silver telluride, or iodine, loaded into polyethylene tubes, and inserted into the phantom. A Hologic 3Dimensions mammography system was used to image the phantom using a tungsten anode and a 70-µm detector. High-energy (HE) and low-energy (LE) images were acquired, from which a weighted logarithmic subtraction was performed. Various pairs of HE and LE images were acquired, HE images were acquired at 49 kV using a copper filter and 90 mAs. The LE images were acquired at every kV from 20 to 39 kV using a silver filter and 100 mAs. Three sets of images were collected at each kV and analyzed using ImageJ. The contrast-to-noise ratios were calculated according to a previous study and a one-way ANOVA with Tukey’s multiple comparisons test was used to determine statistical significance.^32^

#### Computed Tomography

A MI Labs microCT scanner was used to measure the attenuation of the Ag_2_S-NP and Ag_2_Te-NP. The phantom was prepared in triplicate with agents in the following concentrations: 0, 0.5, 1, 2, 4, 6, 8, and 10 mg/mL. These samples were loaded in 200 µL tubes, placed in a rack, and covered in parafilm. The phantom computed tomography (CT) scan was performed with a tube voltage of 55 kV and isotropic 100-micron voxels. Osirix MD software and a one-way ANOVA with Tukey’s multiple comparisons test were used to determine statistical significance.

#### Catalase-Mimetic Activity Assay

The catalase-mimetic activity protocol was adapted from a previous paper.^27^ In brief, an Amplex red hydrogen peroxide/peroxidase assay kit was used. CeNP of various concentrations (i.e 0, 0.01, 0.025, 0.05, 0.1, 0.2, 0.3, 0.5, and 1 mg/mL) were prepared in the reaction buffer. 50 µL of each concentration was added to a 96-well plate. Following this, 50 µL of 40 uM hydrogen peroxide was added to the wells and was incubated for 20 minutes. Finally, 50 µL of working solution was added to each well and the reaction proceeded for another 30 minutes at room temperature. The absorbance was then read at 560 nm.

#### SOD-Mimetic Activity Assay

The SOD activity was measured based on a previously established protocol using a SOD colorimetric activity kit.^27^ CeNP were diluted using PBS to the concentrations of 0.003, 0.007, 0.01, 0.025, 0.05, 0.1, 0.2, 0.3, 0.5, and 1 mg/mL. 10 µL of each concentration was added to a 96-well plate. The substrate and xanthine oxidase were added to each of the wells and the resulting mixtures were incubated at room temperature for 20 minutes. The absorbance was then read at 450 nm.

#### *In Vitro* Cerium Oxide Protection Against ROS

The protocol below was adapted from a previous paper.^28^ In this assay, 20,000 C2BBe1 or BJ5ta cells in 200 µL of cell culture media were added to each well of a 96-well plate and were then incubated for 24 hours to acclimate. The cells were then incubated with various concentrations of CeNP, hydrogen peroxide, or both for 30 minutes. Additionally, a positive control of L-glutathione with cell media was added. The amount of hydrogen peroxide was 0.075% and 0.0125% for C2BBe1 and BJ5ta cells, respectively. After 30 minutes, the culture media from the wells was discarded, and the cells were washed with PBS. New cell culture media was added (200 µL) and incubated again for 24 hours. A MTS assay was used to measure cell viability. The percentage of viability was calculated compared to the control. A one-way ANOVA with Tukey’s multiple comparisons test was used to determine statistical significance.

#### *In Vitro* Catalytic Activity Assessment

A colorimetric assay using 3,3’,5,5’-tetramethylbenzidine (TMB) as a probe was used to measure the catalytic activity of IONP following a previously established protocol.^55^ In brief, 40 µL of the sample (sodium acetate buffer, as a control, and two IONP conditions) and 4 µL of the TMB assay were added to 922 µL of 0.1 M sodium acetate buffer and mixed thoroughly. The absorbance was then recorded using the Genesys 150 UV-visible spectrophotometer at 652 nm. Following this, 34 µL of H_2_O_2_ was added to the control and one of the IONP conditions. Catalytic activity was then monitored at 652 nm every minute for five minutes total.

#### Oral Biofilm Model

A previously established protocol was used.^26,29,68^ In brief, biofilms were formed on vertically suspended saliva-coated hydroxyapatite (sHA) disks (surface area: 2.7 +/- 0.2 cm^2^) in 24 well plates. Filter-sterilized saliva was used to coat each disk for 1 hour at 37 °C. *Streptococcus mutans* UA159 (ATCC 700610) was grown in ultra-filtered (10 kDa, cutoff) tryptone-yeast extract broth (UFTYE; 2.5% tryptone and 1.5% yeast extract, pH 7.0) containing 1% (wt/vol) glucose at 37°C and 5% CO_2_. The bacteria were grown until midexponential phase was reached. Approximately 2 x 10^5^ colony forming units (CFU) of *S. mutans* per mL in UFTYE culture medium containing 1% (wt/vol) sucrose was inoculated onto these sHA disks at 37 °C.

#### *In Vitro* Antibiofilm Effect of IONP

Topical treatment of IONP (1 mg of Fe/mL) or the vehicle control containing sodium acetate buffer (0.1 M NaOAc, pH 4.5) was performed for 10 minutes at 0, 6, 19, 29, and 43 hours on the sHA disks and biofilms. Following treatment, the biofilms were washed thrice with a sterile saline solution and the sHA disks were transferred to culture media. The culture media was changed at 19 and 29 hours. After 43 hours, the biofilms were exposed to 2.8 mL of 1% H_2_O_2_ for 5 minutes. The biofilms were then removed using a spatula and homogenized using batch sonication followed by probe sonication at an output of 7 W for 30 seconds. The homogenized solution was serially diluted and plated onto blood agar plates using EddyJet Spiral Plater. The number of viable cells in each biofilm was determined by counting CFUs and a one-way ANOVA with Tukey’s multiple comparisons test was used to determine statistical significance.

## Statistical Analysis

All experiments were performed in triplicates and labeled below as mean ± standard error mean (SEM) unless otherwise specified. The statistical tests described in the method section were performed using GraphPad Prism 10 software.

## Acknowledgments

The authors would like to acknowledge the staff of the Electron Microscopy Resource Lab, especially S. Molugu, B. Zhou, and I. Martynyuk, for their assistance in acquiring transmission electron micrographs. We also thank E. Blankemeyer for his assistance with micro-CT imaging. Additionally, we thank D. Burney for his assistance with ICP-OES. Finally, we thank the Materials Characterization Core at Drexel University, especially D. Barbash, for his assistance with XPS experiments.

## Author Contributions

The project was designed by A.K., H.K., and D.P.C. The experiments for data acquisition performed by A.K., M.G., H.H., K.L.L.R., N.P., K.M., D.N.R.B., P.S.M., A.R.H., P.S., Z.X., H.K., and D.P.C.. Data analysis was performed by A.K., M.G, H.H., K.L.L.R., N.P., A.D.A.M., D.I., H.K., and D.P.C.. A.K and D.P.C. wrote the manuscript. All authors were involved in the approval of the manuscript.

## Data Availability Statement

The authors confirm that the data supporting the findings of this study are available within Zenodo at https://doi.org/10.5281/zenodo.17592975.

