## Supplemental File for "Scalable flow synthesis of ultrasmall inorganic nanoparticles for biomedical applications *via* a confined impinging jet mixer"

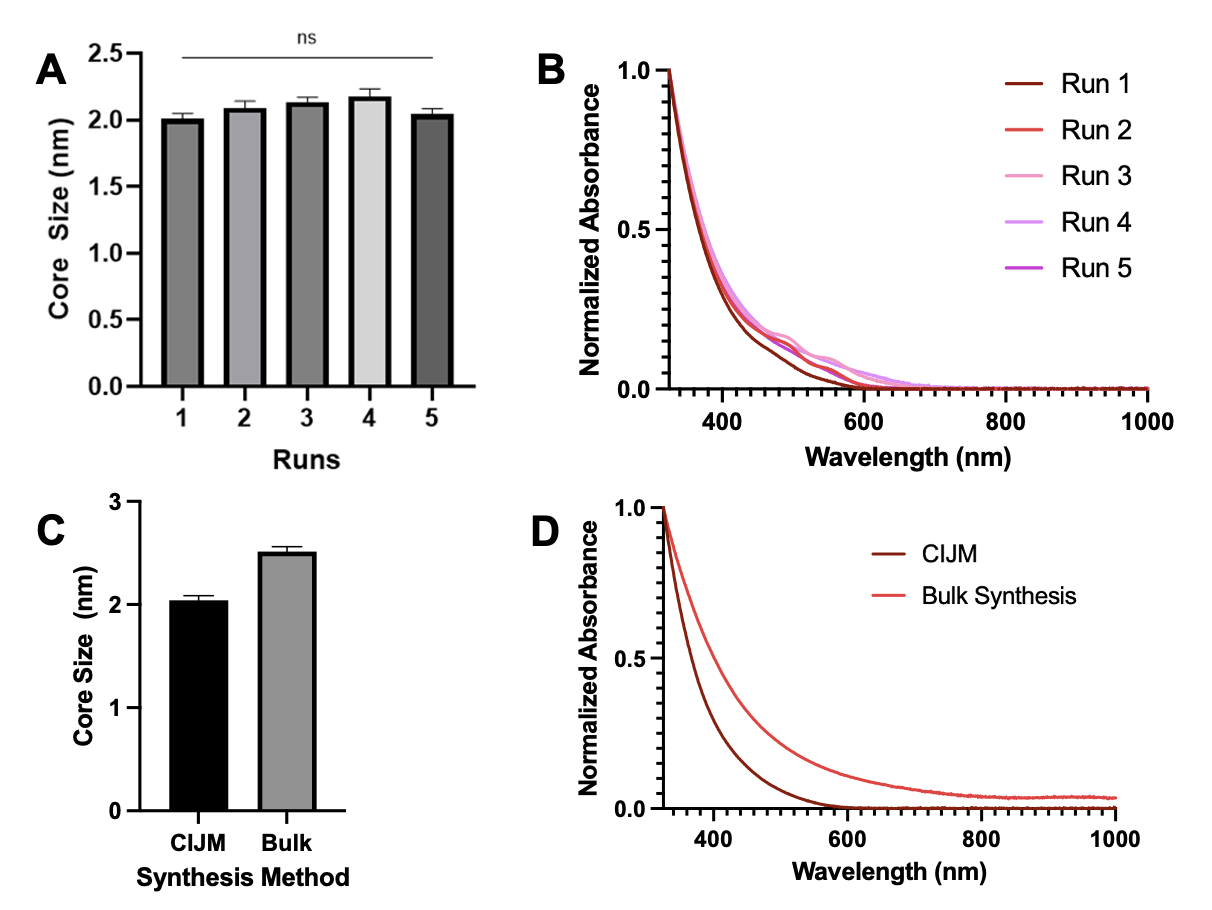

**Figure S1. (A)** Core sizes of different batches of Ag_2_Te-NP. **(B)** UV/vis absorption spectra of the different batches of Ag_2_Te-NP. **(C)** Comparison of core sizes in the CIJM-synthesized and bulk synthesized Ag_2_Te-NP. **(D)** UV/vis absorption spectra of the CIJM-synthesized and bulk synthesized Ag_2_Te-NP. ns = non-significant

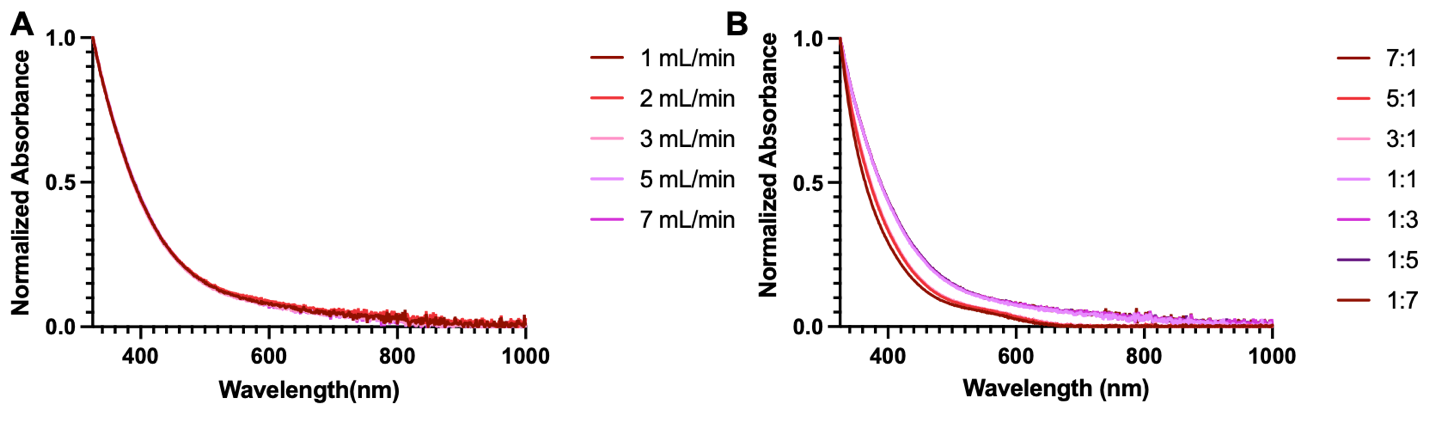

**Figure S2.**  **(A)** UV/vis absorption spectra of Ag_2_S-NP synthesized with differing flow rates (1, 2, 3, 5 and 7 mL/min). **(B)** UV/vis absorption spectra of Ag_2_S-NP synthesized with differing flow rate ratios (7:1, 5:1, 3:1, 1:1, 1:3, 1:5, and 1:7).

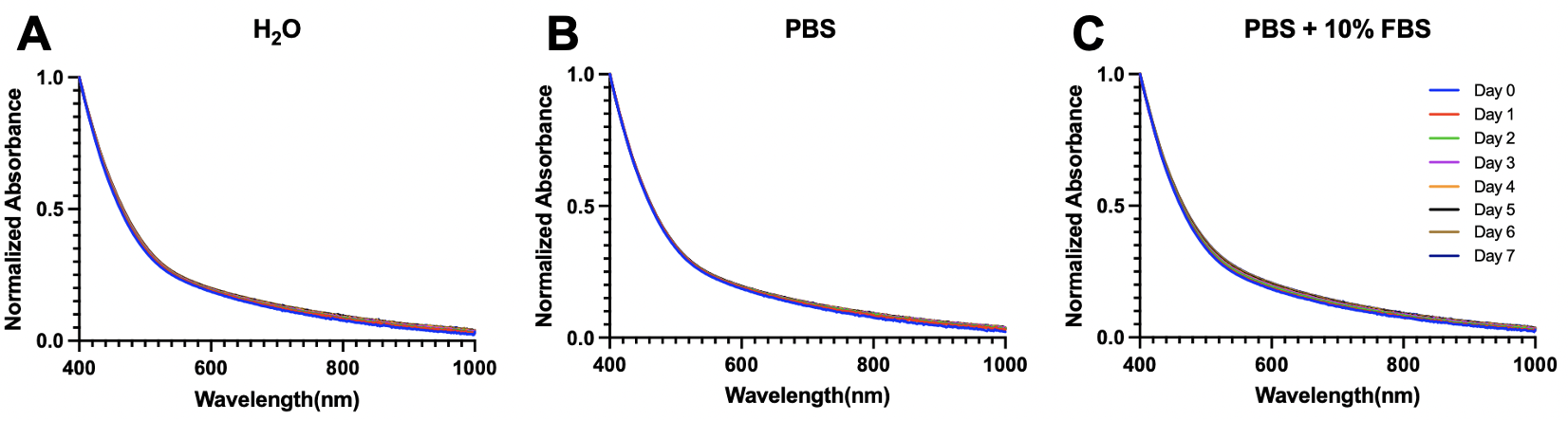

**Figure S3.** UV/vis adsorption spectra of Ag_2_S-NP incubated in **(A)** DI water, **(B)** PBS, and **(C)** PBS + 10% FBS for seven days at 37° C.

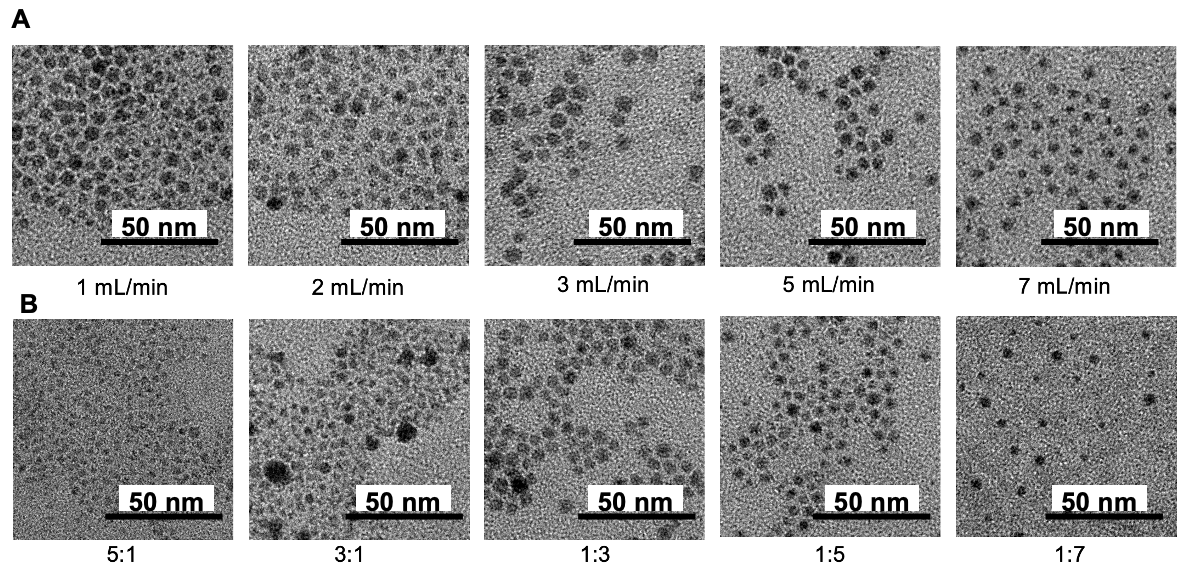

**Figure S4. (A)** Transmission electron micrographs of Ag_2_S-NP synthesized at different flow rates (1, 2, 3, 5 and 7 mL/min). **(B)** TEM of Ag_2_S-NP synthesized at different flow rate ratios (5:1, 3:1, 1:3, 1:5, and 1:7). All scale bars are 50 nm.

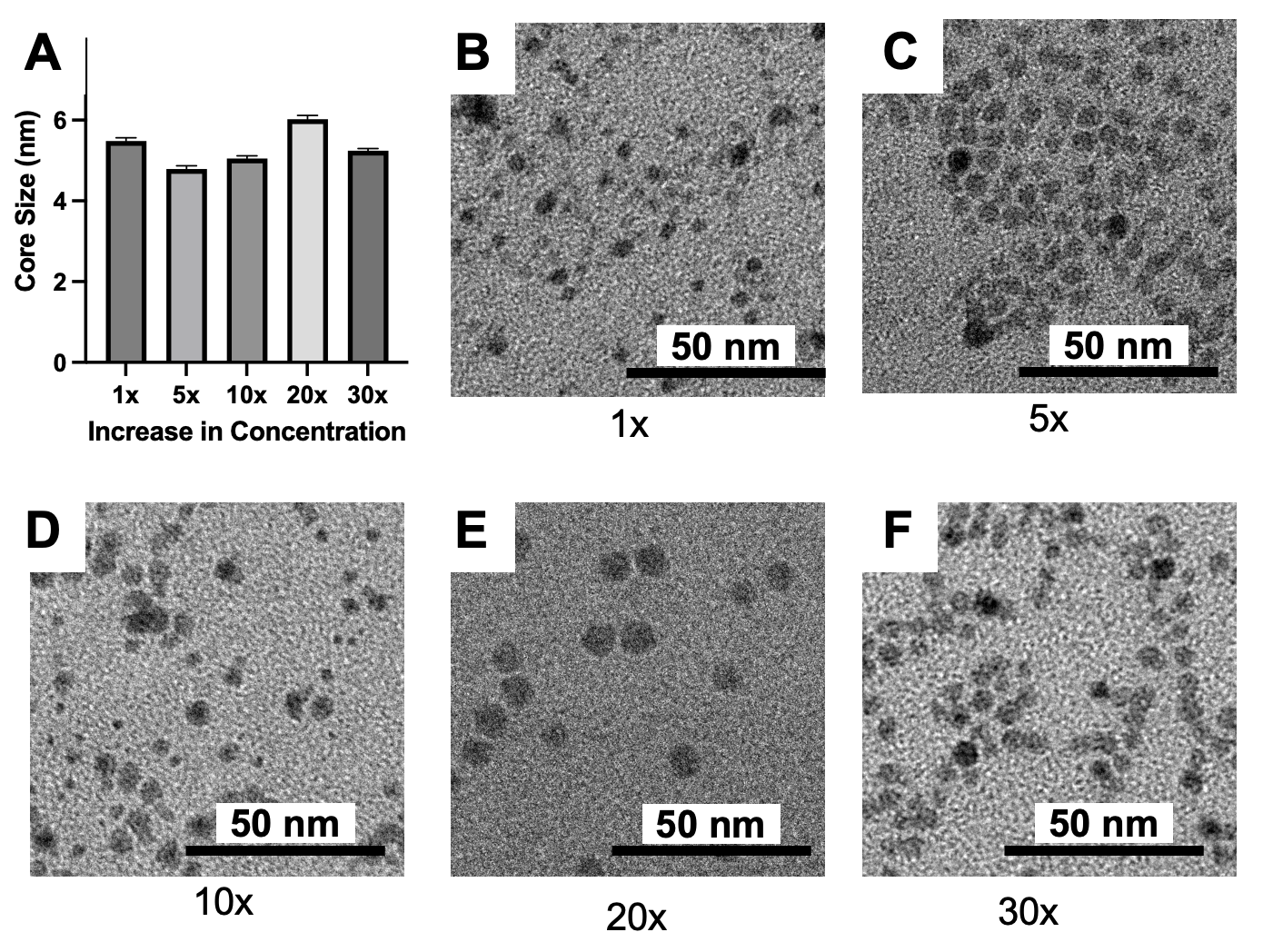

**Figure S5. (A)** Core size of Ag_2_S-NP synthesized at increasing concentrations (1x, 5x, 10x, 20x and 30x). **(B-F)** Transmission electron micrographs of Ag_2_S-NP at increasing concentrations. All scale bars are 50 nm.

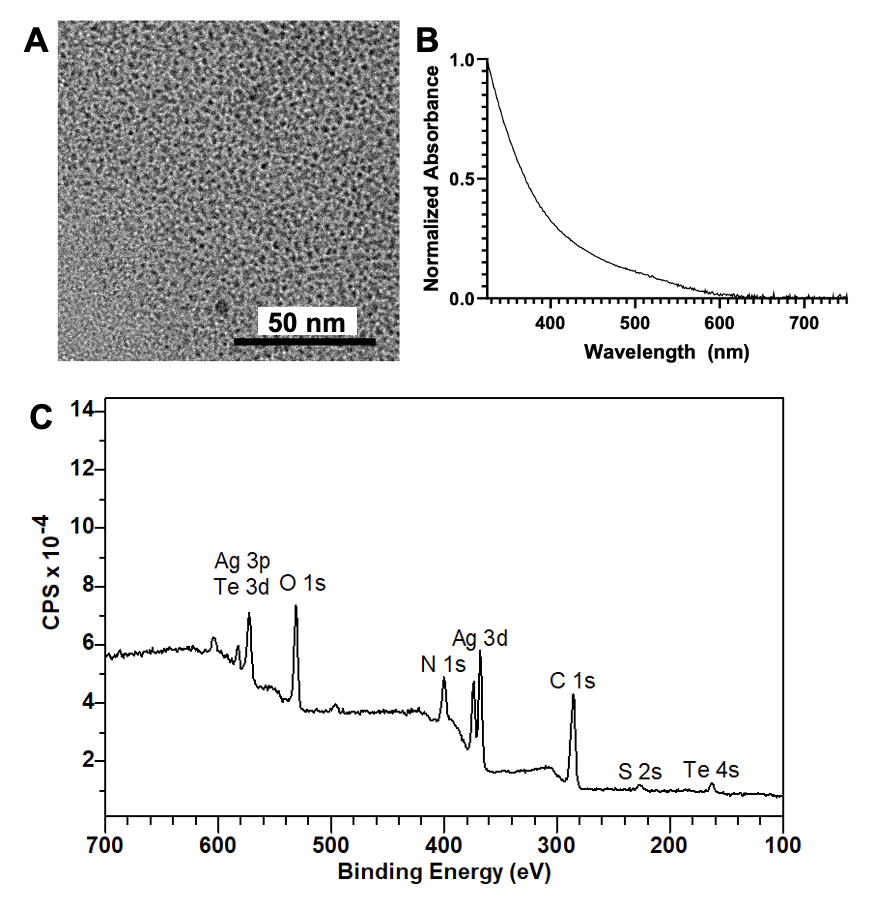

**Figure S6. (A)** Transmission electron micrograph of CIJM-synthesized Ag_2_Te-NP. **(B)** UV/vis absorption spectra of Ag_2_Te-NP. **(C)** XPS of Ag_2_Te-NP. All scale bars are 50 nm.

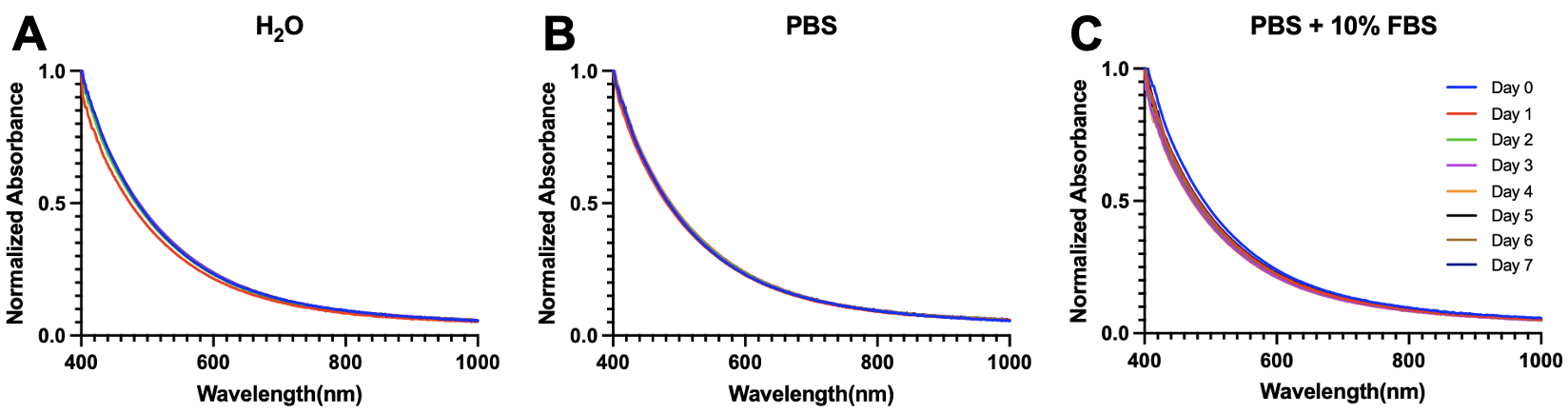

**Figure S7.** UV/vis adsorption spectra of Ag_2_Te-NP incubated in **(A)** DI water, **(B)** PBS, and **(C)** PBS + 10% FBS for seven days at 37° C.

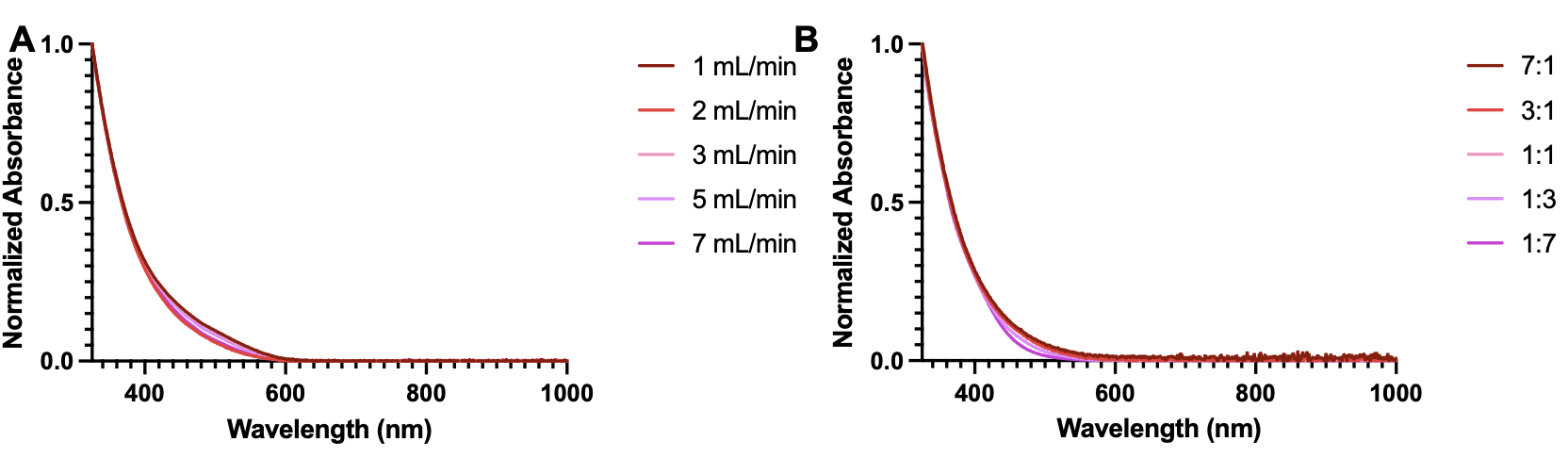

**Figure S8. (A)** UV/vis absorption spectra of Ag_2_Te-NP synthesized with different flow rates (1, 2, 3, 5 and 7 mL/min). **(B)** UV/vis absorption spectra of Ag_2_Te-NP synthesized with different flow rate ratios (7:1, 3:1, 1:1, 1:3, and 1:7).

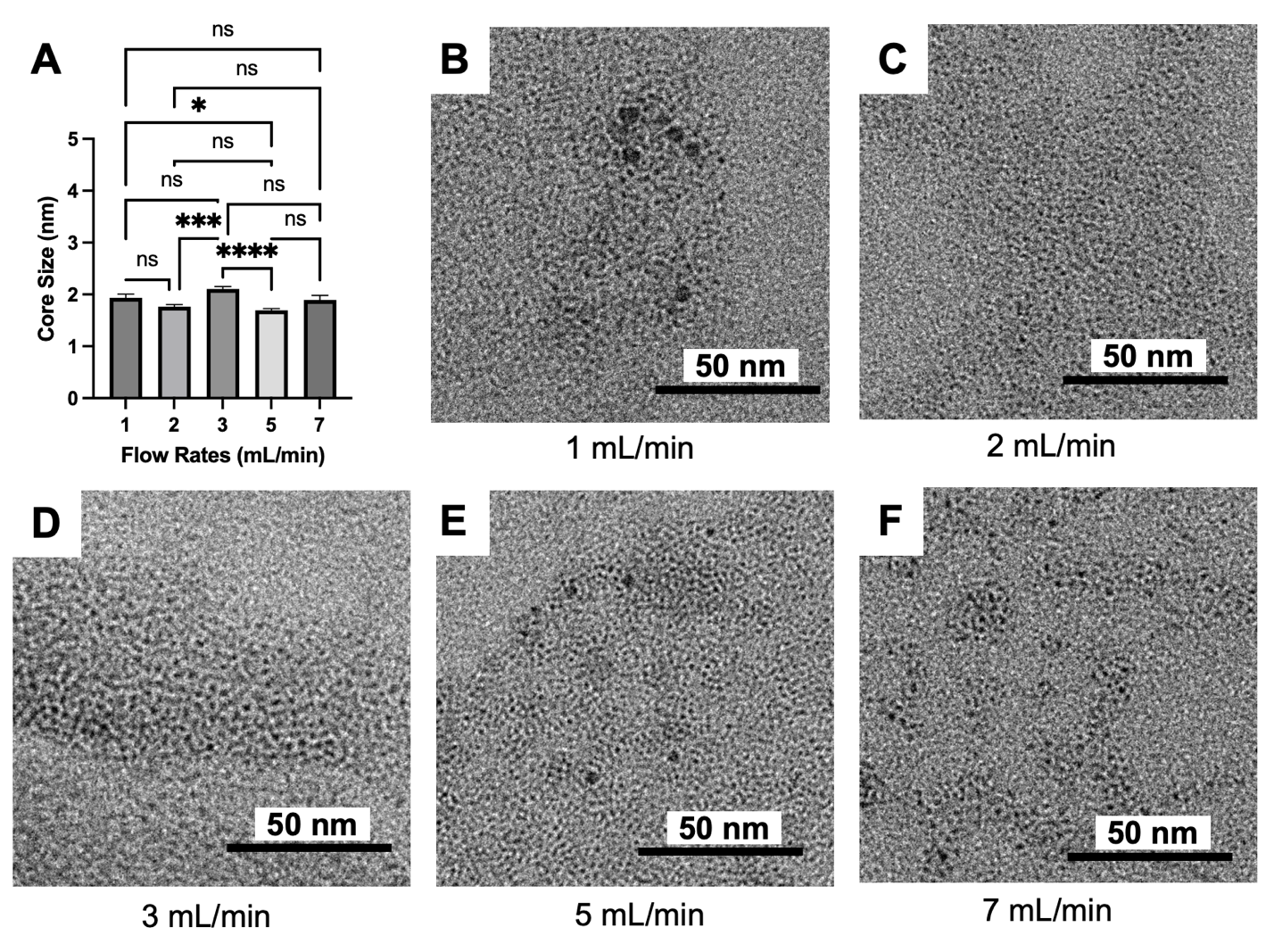

**Figure S9. (A)** Core size of Ag_2_Te-NP synthesized at different flow rates (1, 2, 3, 5 and 7 mL/min). **(B-F)** Transmission electron micrographs of Ag_2_Te-NP at the flow rates noted. All scale bars are 50 nm.

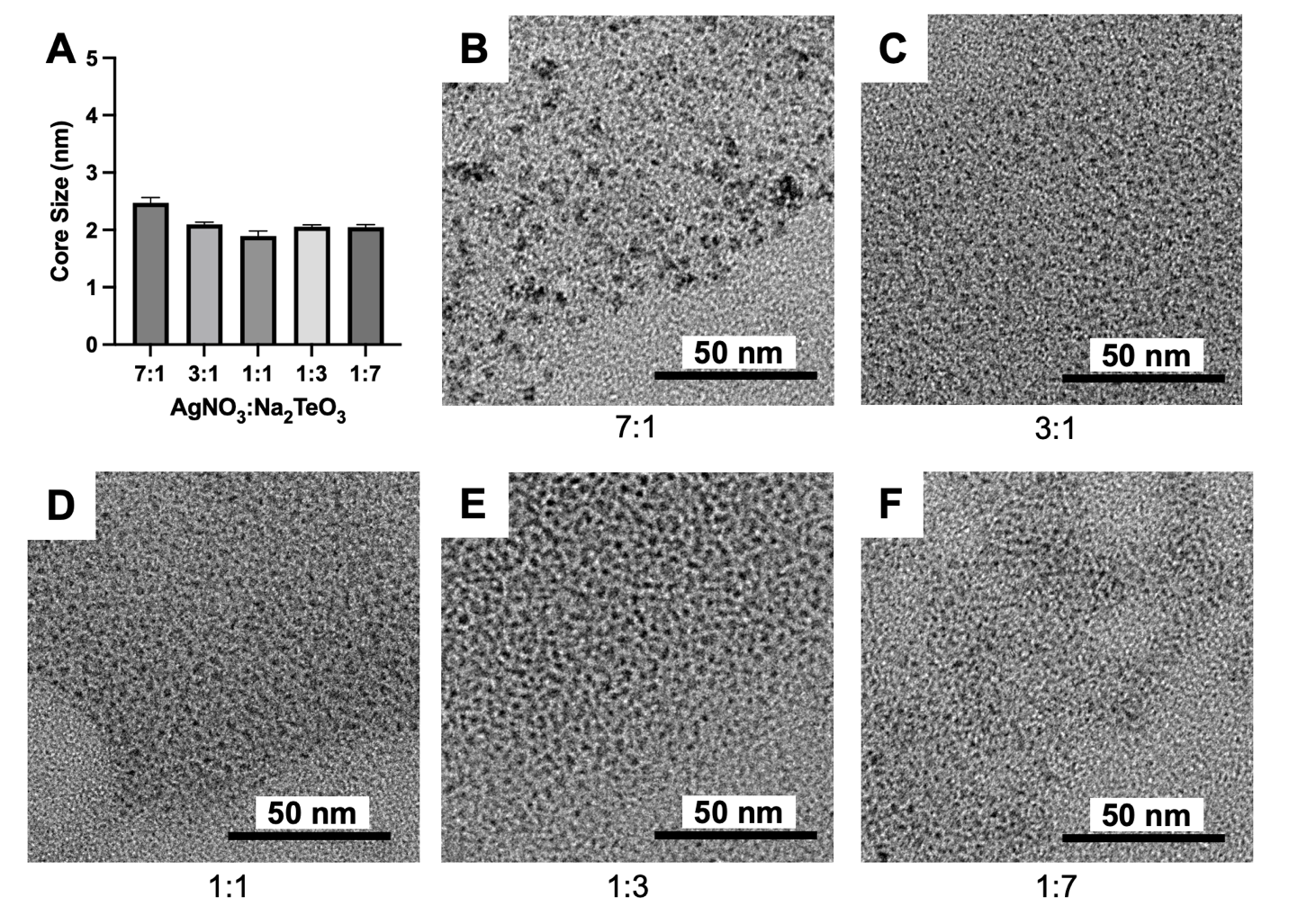

**Figure S10. (A)** Core size of Ag_2_Te-NP synthesized at different flow rate ratios (7:1, 3:1, 1:1, 1:3, and 1:7). **(B-F)** Transmission electron micrographs of Ag_2_Te-NP synthesized at the flow rate ratios noted. Scale bars are 50 nm.

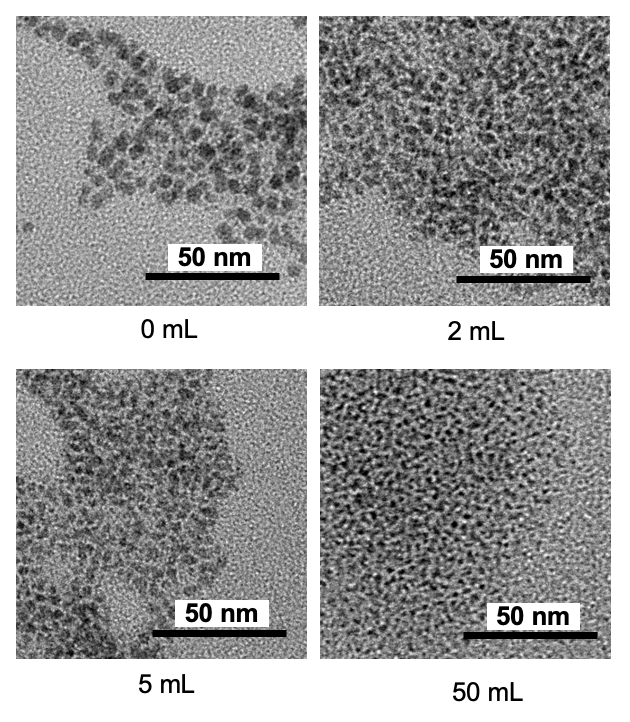

**Figure S11.** Transmission electron micrographs of Ag_2_Te-NP synthesized with the quenching volumes noted. Scale bars are 50 nm.

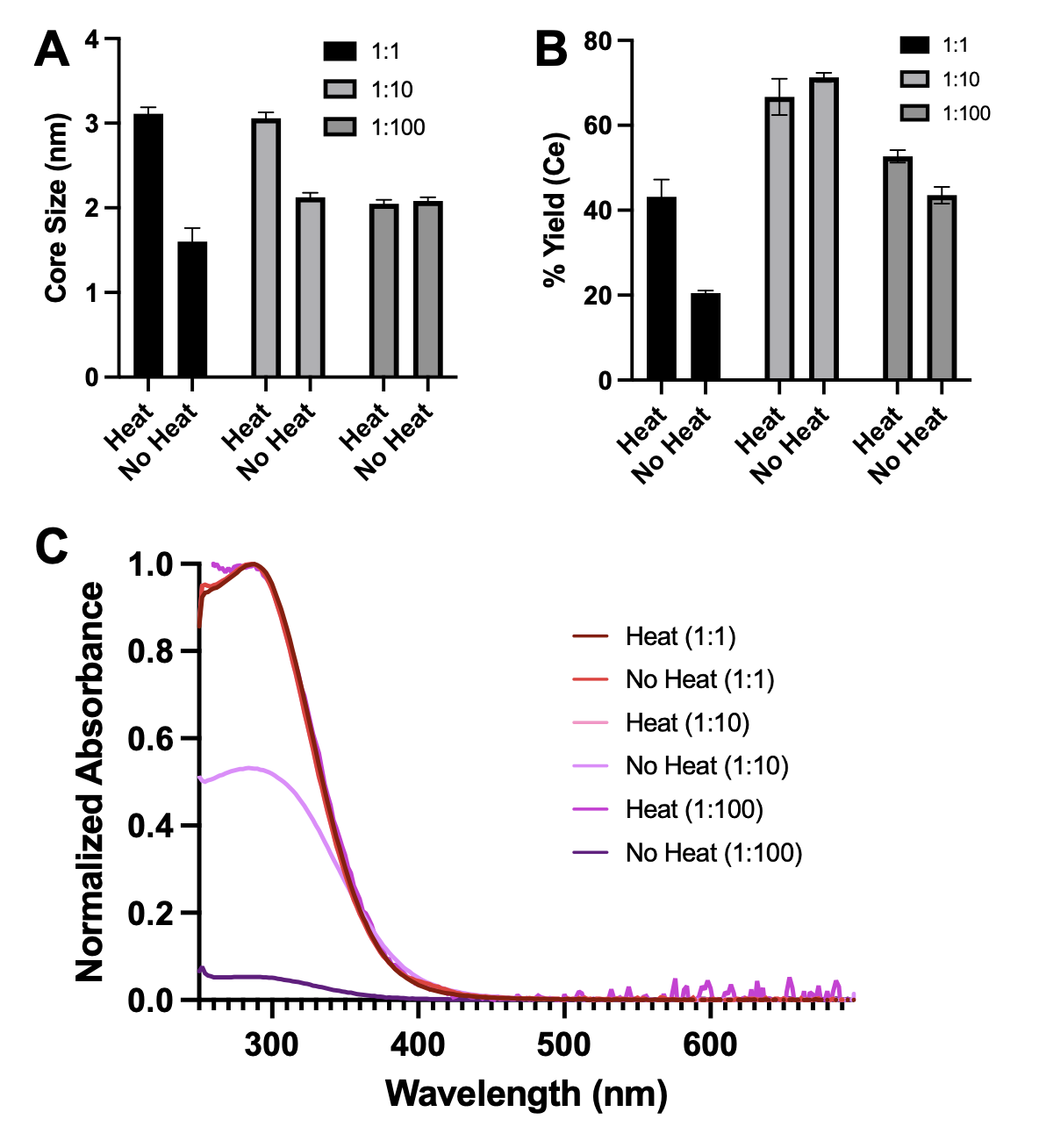

**Figure S12. (A)** Core sizes of CeNP synthesized using different reagent dilutions and using heat (150 °C) or no heat (RT) for the reaction. **(B)** Yield (with respect to cerium) of CeNP synthesized using different reagent dilutions and using heat or no heat. **(C)** UV/vis absorption spectra of the different diluted formulations of CeNP and altering heat and no heat.

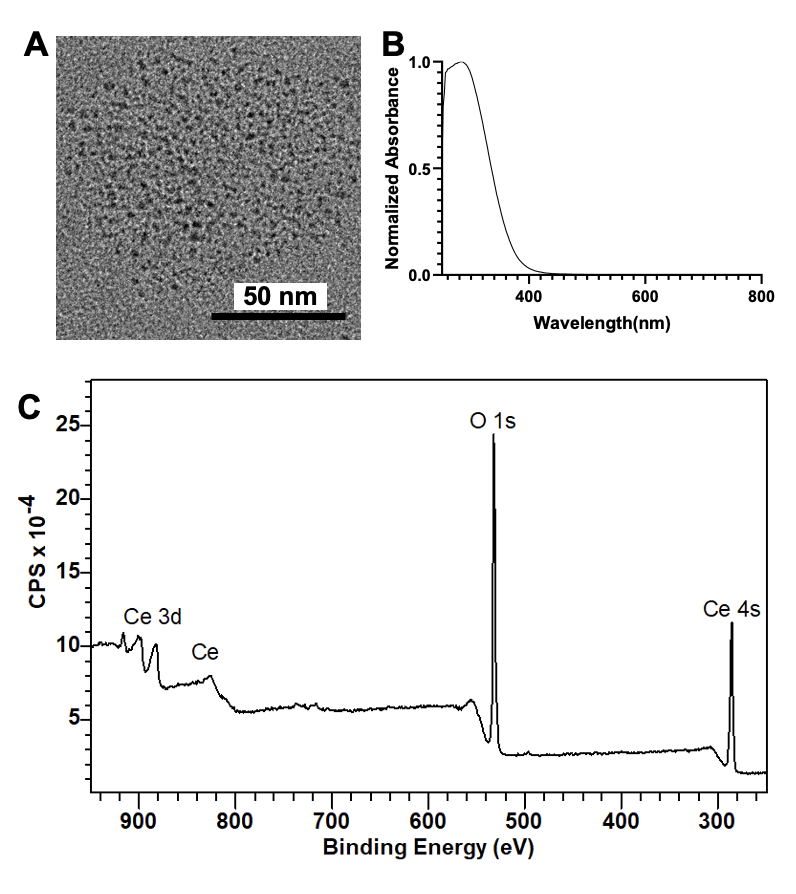

**Figure S13. (A)** Transmission electron micrograph of CeNP. **(B)** UV/vis absorption spectra of CeNP. **(C)** XPS of CeNP.

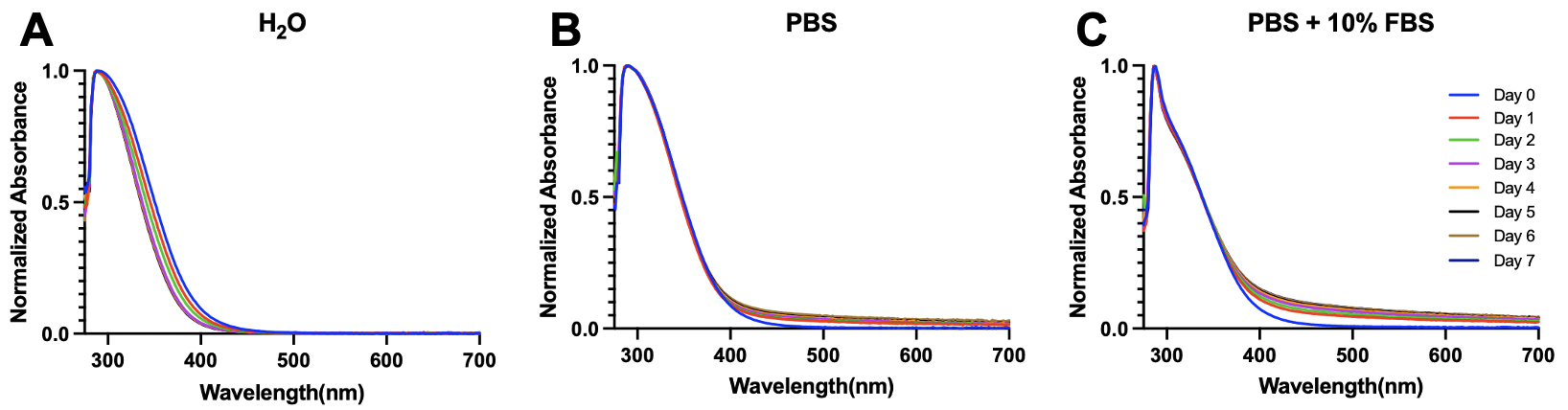

**Figure S14.** UV/vis adsorption spectra of CeNP incubated in **(A)** DI water, **(B)** PBS, and **(C)** PBS + 10% FBS for seven days at 37° C.

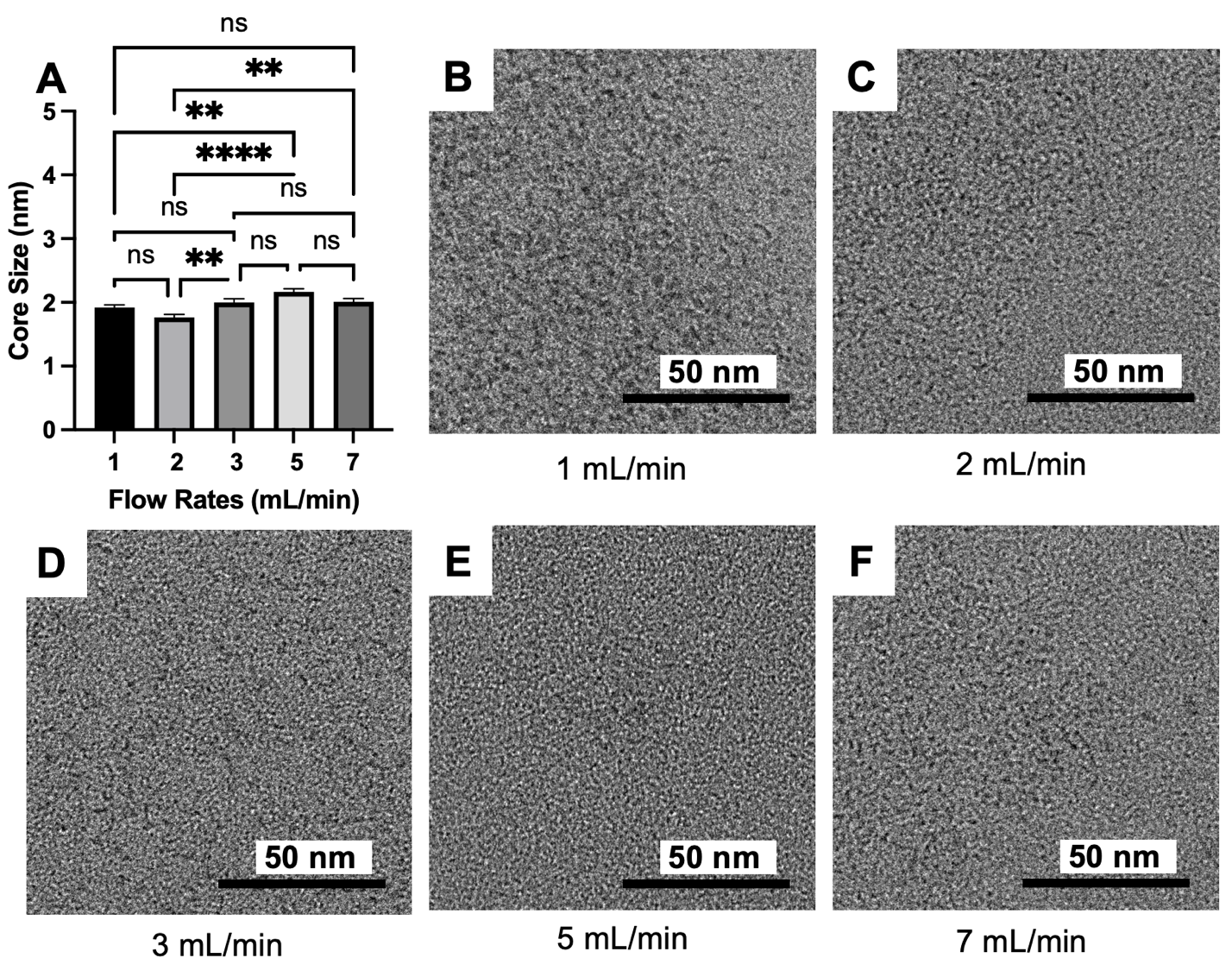

**Figure S15. (A)** Core size of CeNP synthesized at different flow rates (1, 2, 3, 5, and 7 mL/min). **(B-F)** Transmission electron micrographs of CeNP synthesized using the flow rates noted. All scale bars are 50 nm. ns = non-significant, * indicates p≤0.05, ** p≤0.01, *** p≤0.001, **** p≤0.0001.

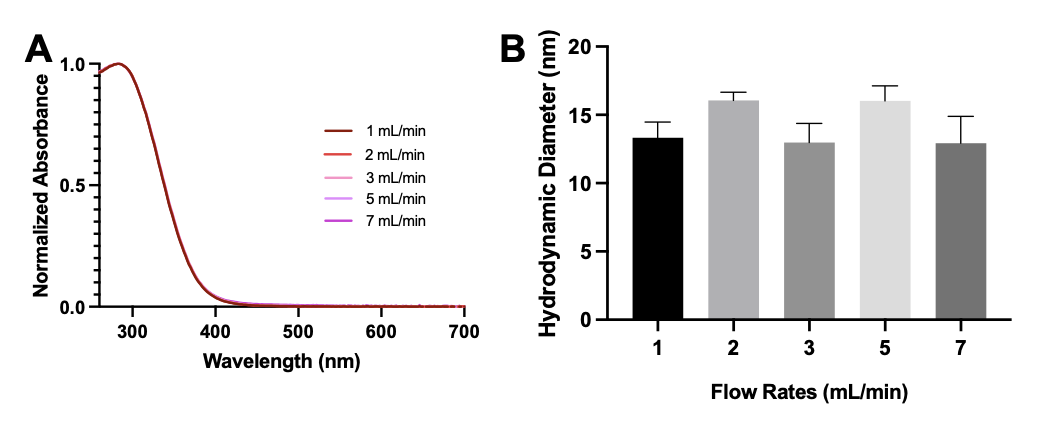

**Figure S16. (A)** UV/vis absorption spectra of CeNP synthesized using the flow rates noted. **(B)** Hydrodynamic diameters of CeNP synthesized using the flow rates noted.

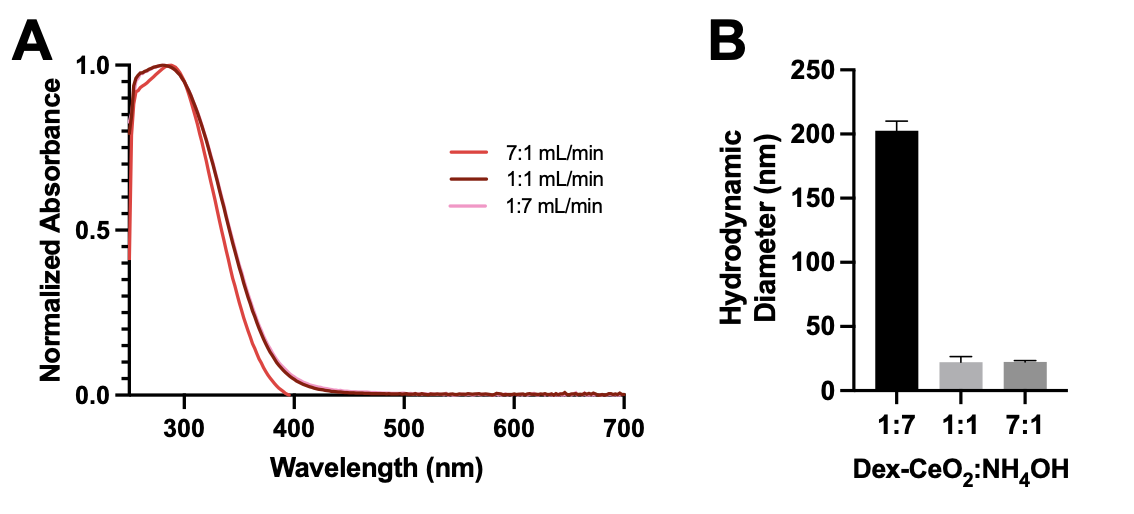

**Figure S17. (A)** UV/vis absorption spectra of CeNP synthesized at the flow rate ratios noted. **(B)** Hydrodynamic diameters of CeNP synthesized using the flow rate ratios noted.

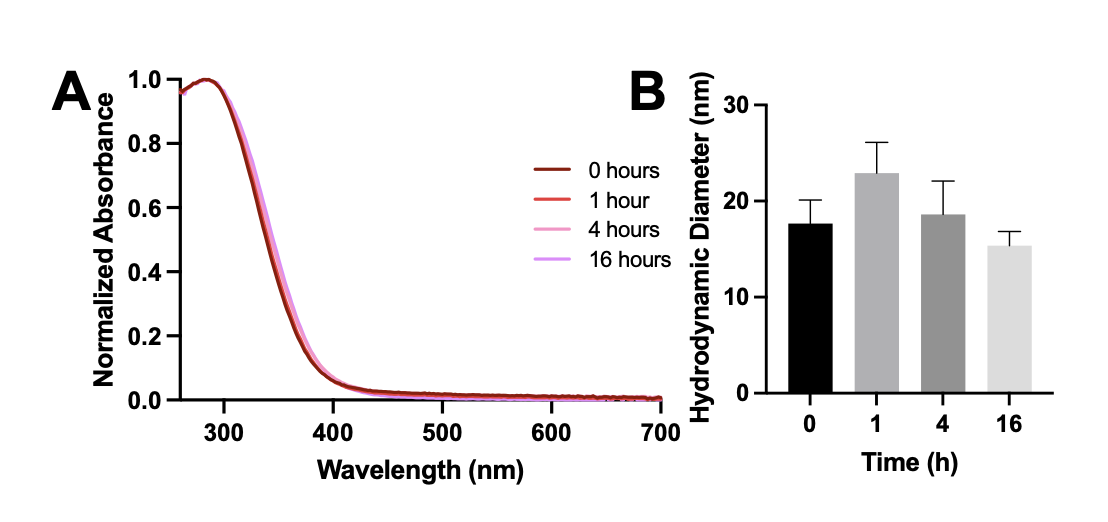

**Figure S18. (A)** UV/vis absorption spectra of CeNP synthesized using the mixing times noted. **(B)** Hydrodynamic diameters of CeNP synthesized using the mixing times noted.

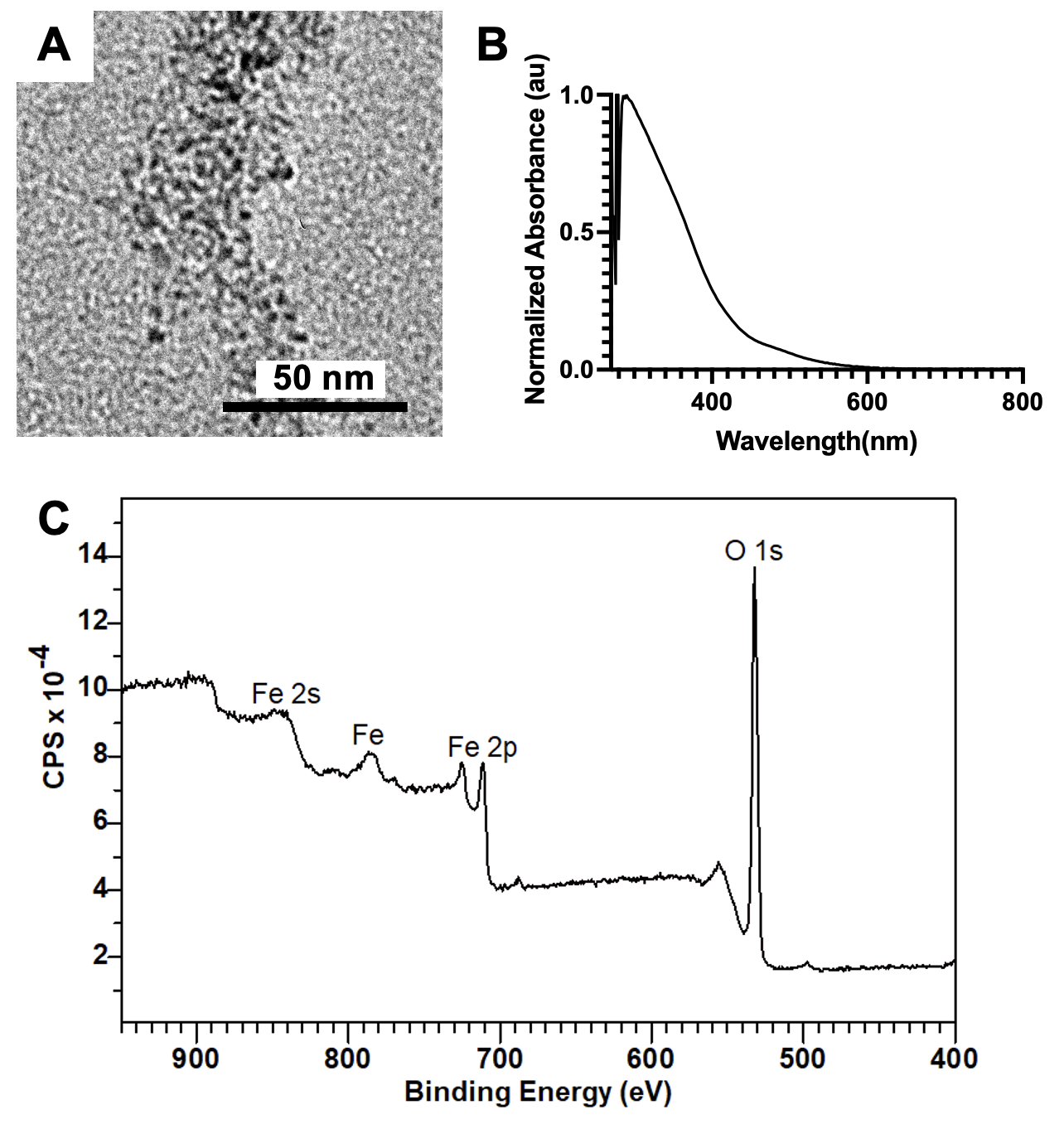

**Figure S19. (A)** Transmission electron micrograph of IONP. **(B)** UV/vis absorption spectrum of IONP. **(C)** XPS of IONP. The scale bar represents 50 nm.

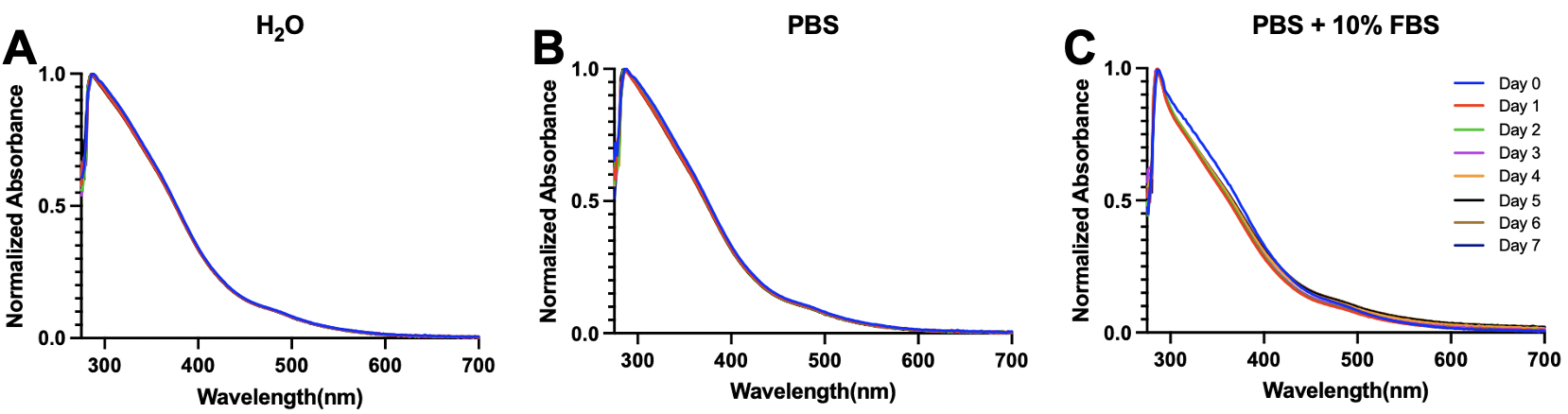

**Figure S20.** UV/vis adsorption spectra of IONP incubated in **(A)** DI water, **(B)** PBS, and **(C)** PBS + 10% FBS for seven days at 37° C.

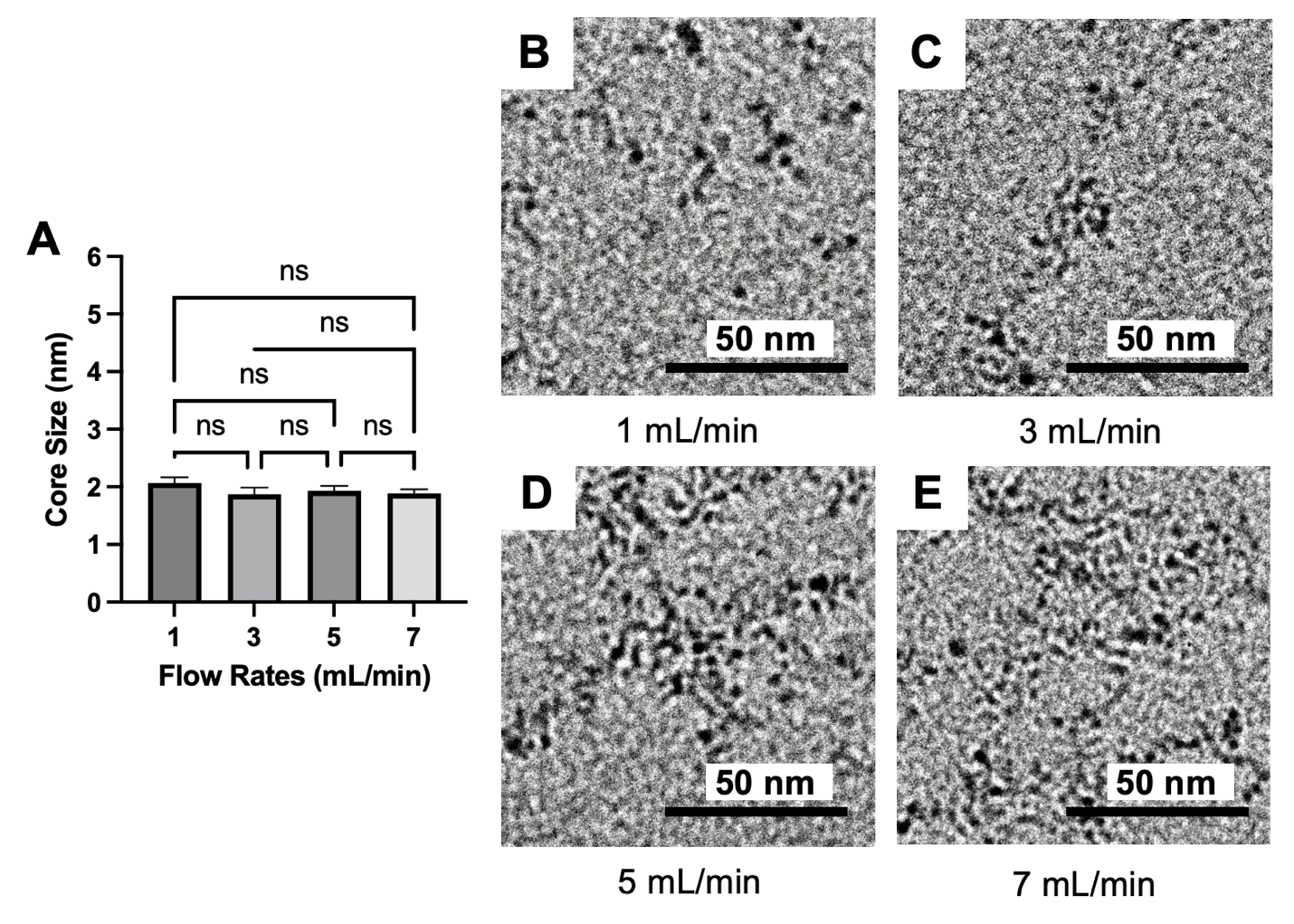

**Figure S21. (A)** Core size of IONP synthesized at different flow rates (1, 3, 5 and 7 mL/min). **(B-E)** Transmission electron micrographs of the IONP synthesized using the flow rates noted. All scale bars are 50 nm. ns = non-significant.

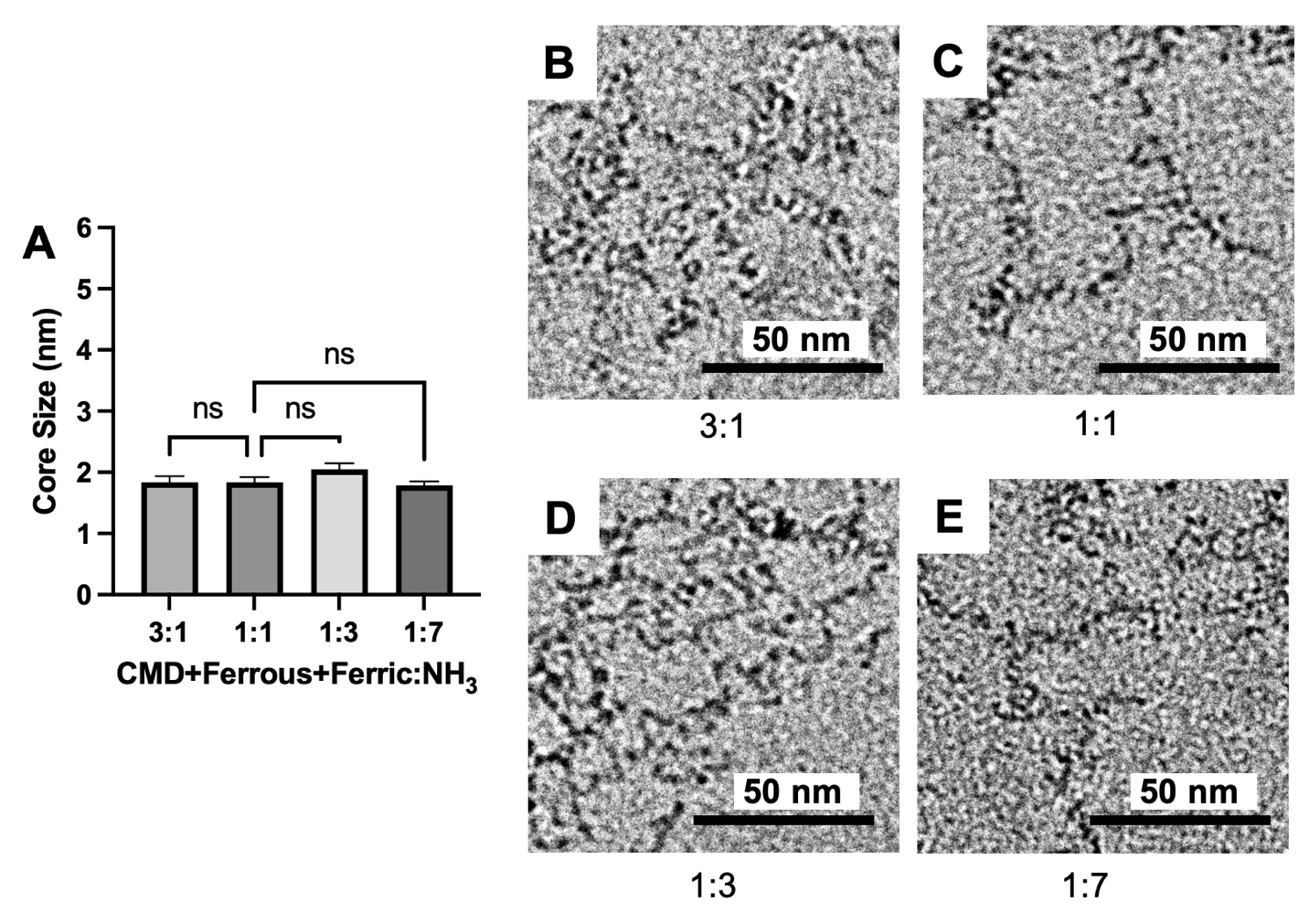

**Figure S22. (A)** Core sizes of IONP synthesized using different flow rate ratios. **(B-E)** Transmission electron micrographs of IONP synthesized the flow rate ratios noted. All scale bars are 50 nm. ns = non-significant.

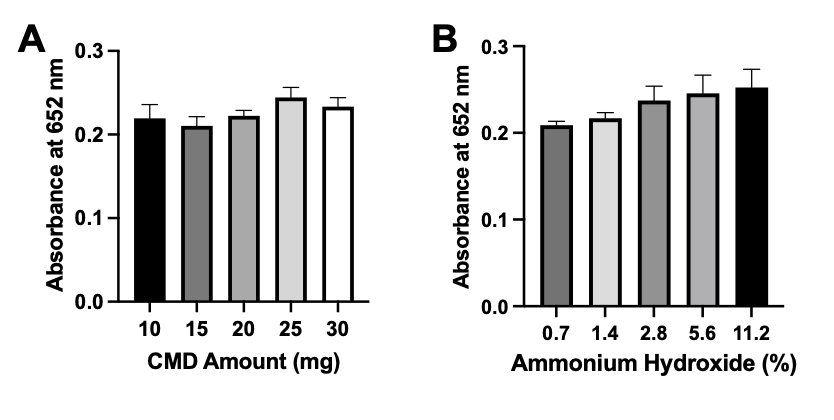

**Figure S23. (A)** Catalytic activity of IONP synthesized using the amounts of CMD noted. **(B)** Catalytic activity of IONP synthesized using the concentrations of ammonia hydroxide noted.

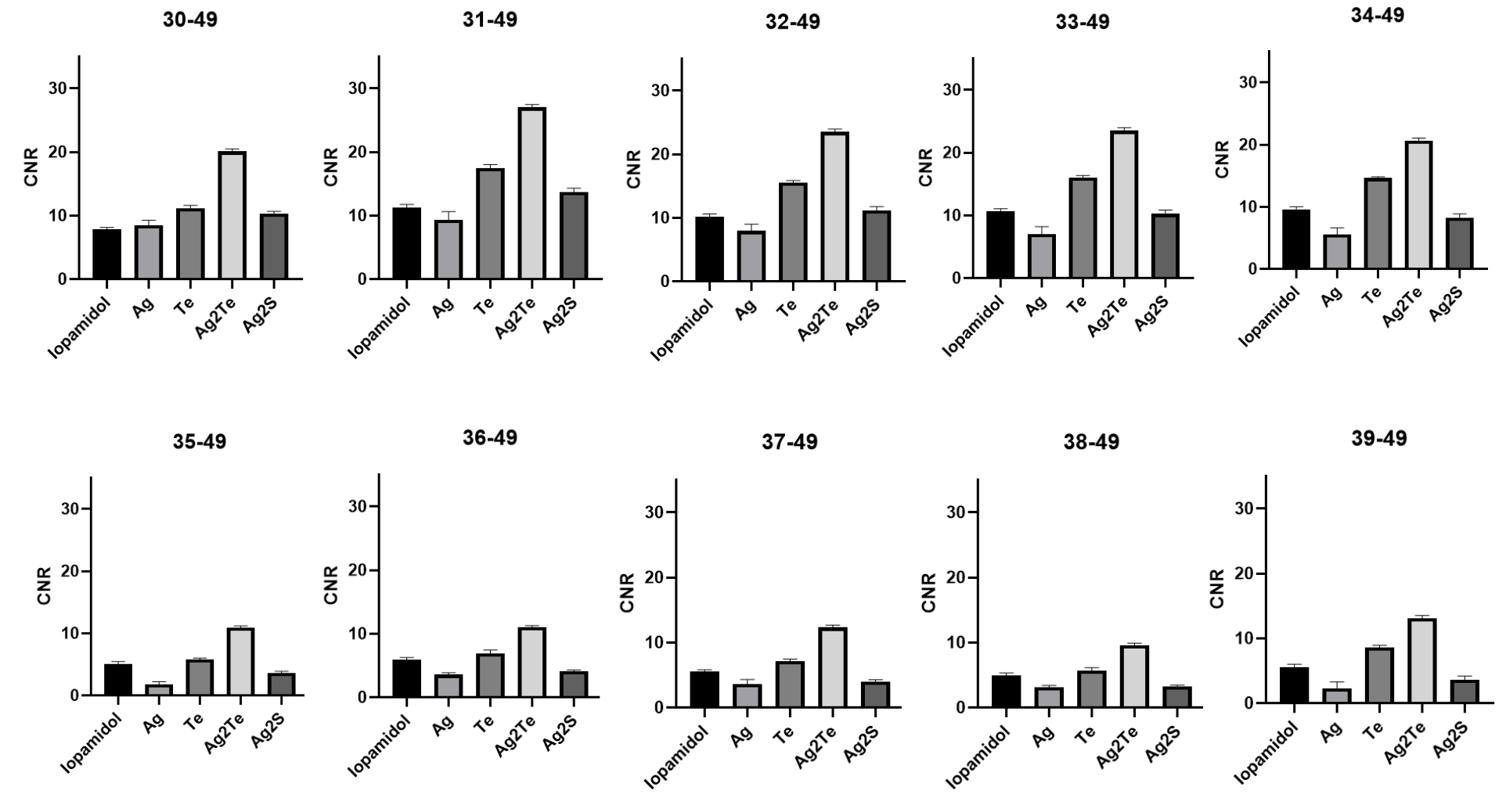

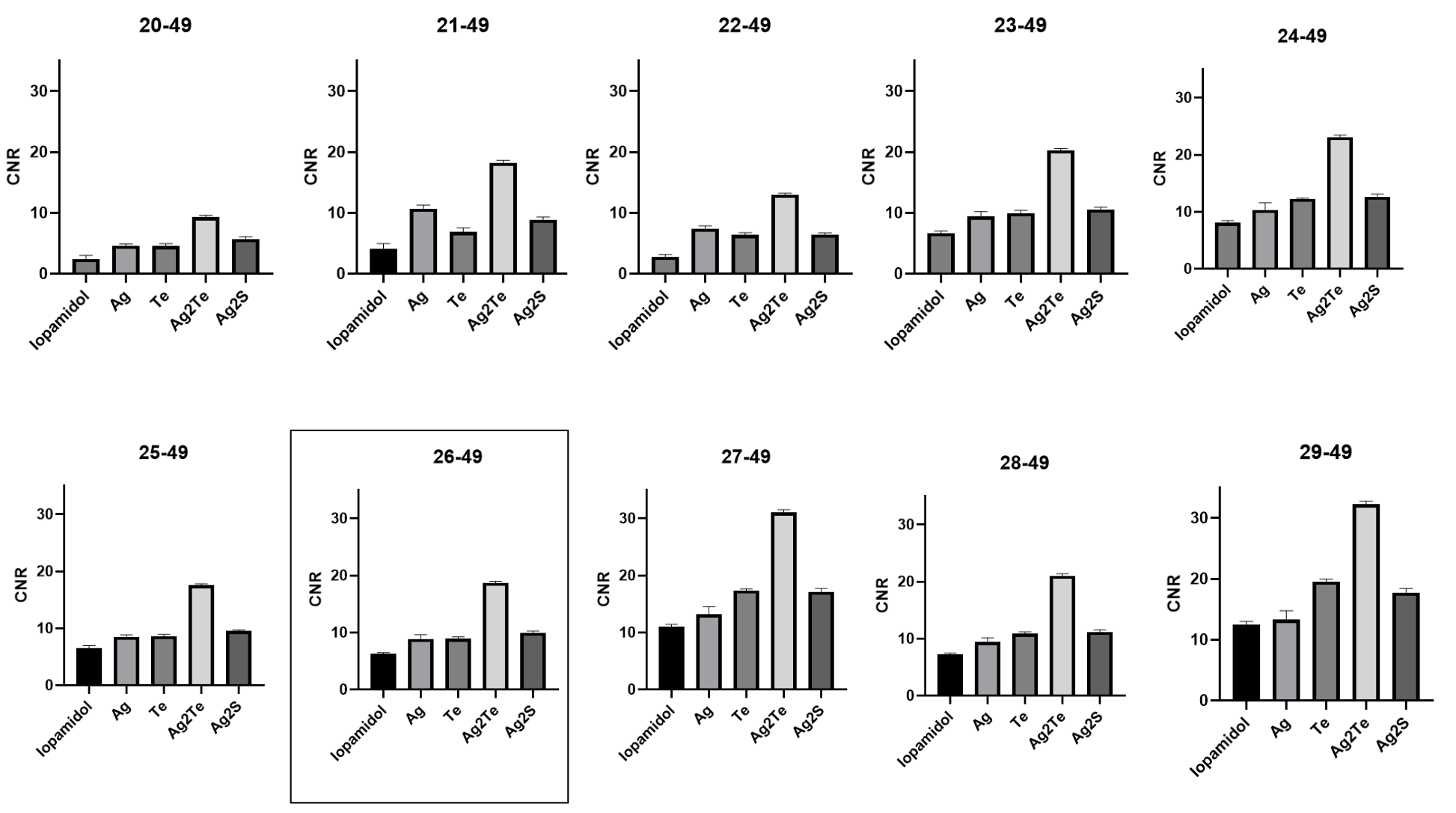

**Figure S24.** CNR in contrast-enhanced mammography images acquired at various low energy-high energy pairs (LE-HE). The samples measured are iopamidol (an iodine-based contrast agent), silver nitrate (noted as Ag), sodium tellurite (noted as Te), Ag_2_Te-NP (noted as Ag_2_Te), and Ag_2_S-NP (noted as Ag_2_S). The boxed pair indicates what was chosen for the main text.

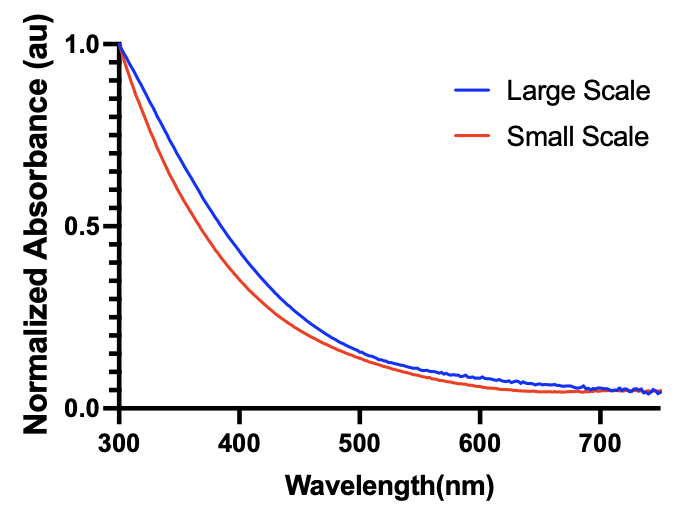

**Figure S25.** UV/vis absorption spectra of Ag_2_S-NP synthesized on large and small scales.
